# Full-length single-cell BCR sequencing paired with RNA sequencing reveals convergent responses to vaccination

**DOI:** 10.1101/2023.05.23.541927

**Authors:** Duncan M. Morgan, Yiming Zhang, Jin-Hwan Kim, MaryAnn Murillo, Suddham Singh, Jakob Loschko, Naveen Surendran, Sarita U. Patil, Isis Kanevsky, Laurent Chorro, J. Christopher Love

**Affiliations:** Koch Institute for Integrative Cancer Research, MIT, Cambridge, Massachusetts, USA; Department of Chemical Engineering, MIT, Cambridge, Massachusetts, USA; Department of Biological Engineering, MIT, Cambridge, Massachusetts, USA; Vaccine Research and Development Pfizer Inc., Pearl River, New York, USA; Center for Immunology and Inflammatory Diseases, Massachusetts General Hospital, Harvard Medical School, Boston, MA

## Abstract

Single-cell RNA sequencing can to resolve transcriptional features from large numbers of individual immune cells, but techniques capable of resolving the variable regions of B cell receptors (BCR) – defining features that confer antigen specificity to B cells – remain limited, especially from widely-used 3′-barcoded libraries. Here, we report a method that for recovering paired, full-length variable region sequences of the BCRs from 3′-barcoded single-cell whole transcriptome libraries. We first verified this method could produce accurate, full-length BCR sequences. We then applied this method to profile antigen-specific B cell responses elicited against the capsular polysaccharide of *Streptococcus pneumoniae* serotype 3 (ST3) by glycoconjugate vaccines in infant rhesus macaques. Using our method, we defined features of the BCR associated with specificity for the ST3 antigen and showed that these sequence characteristics are present in multiple vaccinated monkeys, indicating a convergent response to vaccination. These results demonstrate the utility of our method to resolve key features of the B cell repertoire and for profiling antigen-specific responses elicited by vaccination.

## Introduction

B cells play a pivotal role in the adaptive immune response through the production of antibodies that can exhibit specificity against a diverse array of pathogens. The diversity of the humoral response stems initially from the genetic recombination and junctional diversification of the heavy and light chain subunits that together comprise the B cell receptor (BCR)^1^. Upon recognition of antigen, naïve B cells enter germinal centers (GCs), where they undergo clonal expansion, somatic hypermutation (SHM), and class-switch recombination (CSR)^2–4^. These processes induce a further diversified and affinity-matured BCR repertoire capable of exerting a diverse array of effector functions, depending on the immunological context at hand. GC B cells can also undergo differentiation into either memory B cells or plasma cells, leading to the proliferation of the humoral response and the establishment of immunological memory^5–7^.

Previous analyses of the BCR repertoire have provided insights into antigen-specific immune response in a wide variety of contexts, including autoimmune diseases, allergies, and vaccination^8–14^. Commonly, variable regions from the BCR heavy chain are analyzed *en masse* using next-generation sequencing^8–14^. While this approach provides substantial throughput, it fails to acquire naturally paired heavy and light chain sequences, which are required to express and functionally interrogate the specificity, affinity, and clonality of the antibodies elicited by an immune response. Most frequently, to obtain paired heavy and light chains, single B cells are sorted into microliter plates, and BCRs are subsequently amplified with nested PCR followed by Sanger sequencing for analysis, limiting overall throughput to ∼100s of cells^15–18^. More recently, strategies based on next-generation sequencing have enabled the acquisition of pairings of heavy/light chains and simultaneous readouts of BCR specificity^19–22^, but these methods remain limited in their ability to simultaneously assess the transcriptomes of single B cells. In addition, these methods remain difficult to apply to sparse samples, such as small populations of antigen-specific cells isolated from the blood or biopsy samples available from human patients.

Single-cell RNA sequencing (scRNA-seq) currently affords the ability to analyze the whole transcriptomes of single cells with substantial throughput and has revolutionized studies of gene expression^23, 24^. While pioneering studies combining the analysis of the BCR repertoire with B cell phenotypes have furthered our knowledge of how BCRs relate to the fates and functions of single B cells^25–28^, many scRNA-seq platforms remain limited in their ability to obtain BCR variable region sequences. Early demonstrations of scRNA-seq based on the isolation of individual cells into microliter plates followed by full-length RNA sequencing enabled the in silico reconstruction of BCR variable region sequences^29, 30^. By their nature, these solutions exhibit reduced throughput compared to massively parallel platforms for scRNA-seq, which utilize short sequence reads to obtain digital counts of gene expression, rather than full-length RNA sequence coverage. Because the BCR variable region is located on the 5′-end of the BCR transcript, sufficient BCR variable region coverage can often be obtained from 5′-barcoded library constructions,^31^ but this constraint poses a significant limitation for 3′-barcoded library constructions, which obtain minimal coverage of the BCR variable region. Reported methods to enable the recovery of BCR variable region sequences from 3′-barcoded libraries have relied on specialized RNA capture reagents (DART-seq)^32^ or required the use of multiple sequencing modalities (i.e. sequencing-by-synthesis plus long-read nanopore sequencing)^33, 34^. These constraints have limited the broad adoption of these techniques based on available resources, total costs, or both.

Here, we present an approach for the recovery and sequencing of paired, full-length variable region sequences compatible with standard methods for generating 3′-barcoded scRNA-seq libraries, including Drop-seq, Seq-Well, and other commercial systems^35–37^. The approach is cost effective, can easily be scaled to large numbers of samples, and can be used to recover BCR sequence information from previously archived samples. We first established the ability of our approach to recover accurate BCR sequences from human peripheral blood mononuclear cells (PBMC). We then used the approach to profile both transcriptional and clonotypic features present among antigen-specific B cells elicited by protein-polysaccharide conjugate vaccines in rhesus macaques.

### Targeted recovery and sequencing of full-length, variable region BCR sequences

We sought to extend a method we previously reported for the recovery of the complementarity-determining region 3 (CDR3) sequences of T cell receptors (TCRs)^38^. While full-length BCR transcripts are captured by 3′-barcoded libraries, the process of random fragmentation used to generate size-defined whole-transcriptome libraries, combined with short sequencing reads (∼50-150nt) used leads to loss of coverage of the variable regions of BCR transcripts (Figure 1A). In our approach, a portion of the 3′-barcoded whole-transcriptome amplification (WTA) product generated from standard scRNA-seq protocols is enriched for BCR sequences by probe-based affinity capture using biotinylated oligonucleotides that target the constant regions of BCR heavy and light chain isotypes (Figure 1B). This product is then reamplified using the same universal primer sites (UPS) as the original WTA reaction. The resulting BCR-enriched product is subsequently modified by primer extension using a set of oligonucleotides comprising a shared 5′ UPS (UPS2) linked to regions specific for the leader (L) or framework 1 (FR1) region of BCR heavy and light chain variable (V) segments (Figure 1C). The product of this primer extension step is finally amplified with primers containing sequencing platform-dependent adapters linked to regions specific for either UPS2 (5′-end of construct) or the original UPS (3′-end of construct) (Figure 1D). The resulting amplicons can then be sequenced using two overlapping reads in opposite directions, using the UPS2 sequence appended during primer extension (5′ to 3′ reads) and custom sequencing primers targeting BCR constant regions (3′ to 5′ reads). In addition, a third read is used to obtain the cellular barcode and UMI appended to the BCR during WTA library preparation. The two BCR reads are assembled *in silico* to reconstruct a full-length, variable region BCR sequence for each molecule, spanning from the V-region primer binding site to the beginning of the constant region and can be matched to corresponding single-cell transcriptomes using the corresponding cellular barcodes.

**Figure 1.**
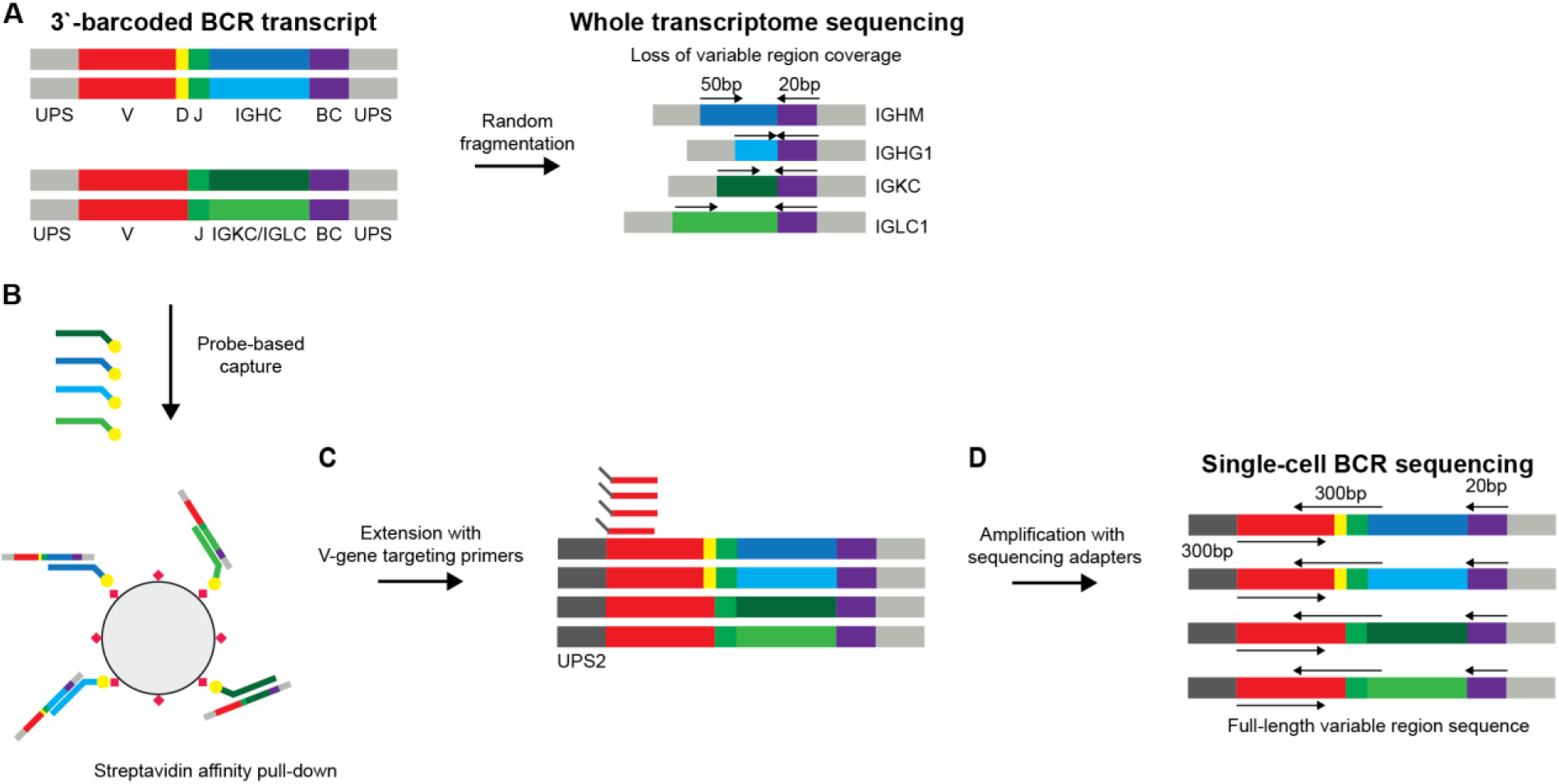
Approach for recovery of full-length variable region BCR sequences from 3′-barcoded sequencing libraries. In whole-transcriptome sequencing, barcoded cDNA is randomly fragmented and amplified to produce size-defined sequencing libraries that are sufficient for transcript enumeration but lead to loss of coverage of BCR variable regions. In our approach, BCR transcripts are enriched from cDNA product using probe-based affinity capture with oligonucleotides targeting the constant regions of BCR isotypes. This enrichment product is then modified by primer extension with hybrid UPS2/V-gene targeting primers and amplified with sequencing adapters to produce a size-defined sequencing library containing the full length of the BCR variable region as well as the single-cell cellular barcode and UMI. During sequencing, one read is used to capture the cellular barcode and UMI, and two overlapping reads in opposite directions are used to capture the full length of the BCR variable region, which can be assembled in silico and matched to single-cell transcriptomes using the corresponding cellular barcode.

### Recovery of BCR sequences from human peripheral blood mononuclear cells

To demonstrate the ability of our method to recover BCR sequences, we performed an initial experiment in which we enriched B cells from human peripheral blood mononuclear cells (PBMC) using magnetic associated cell separation (MACS) and analyzed the resulting suspension of cells with scRNA-seq. In the resulting whole-transcriptome data, we annotated cells according to transcripts expressed and identified two populations of B cells, which correspond to naïve B cells and memory B cells, one small population of plasmablasts, and additional populations of non-B cells (Figure 2A-B).

**Figure 2.**
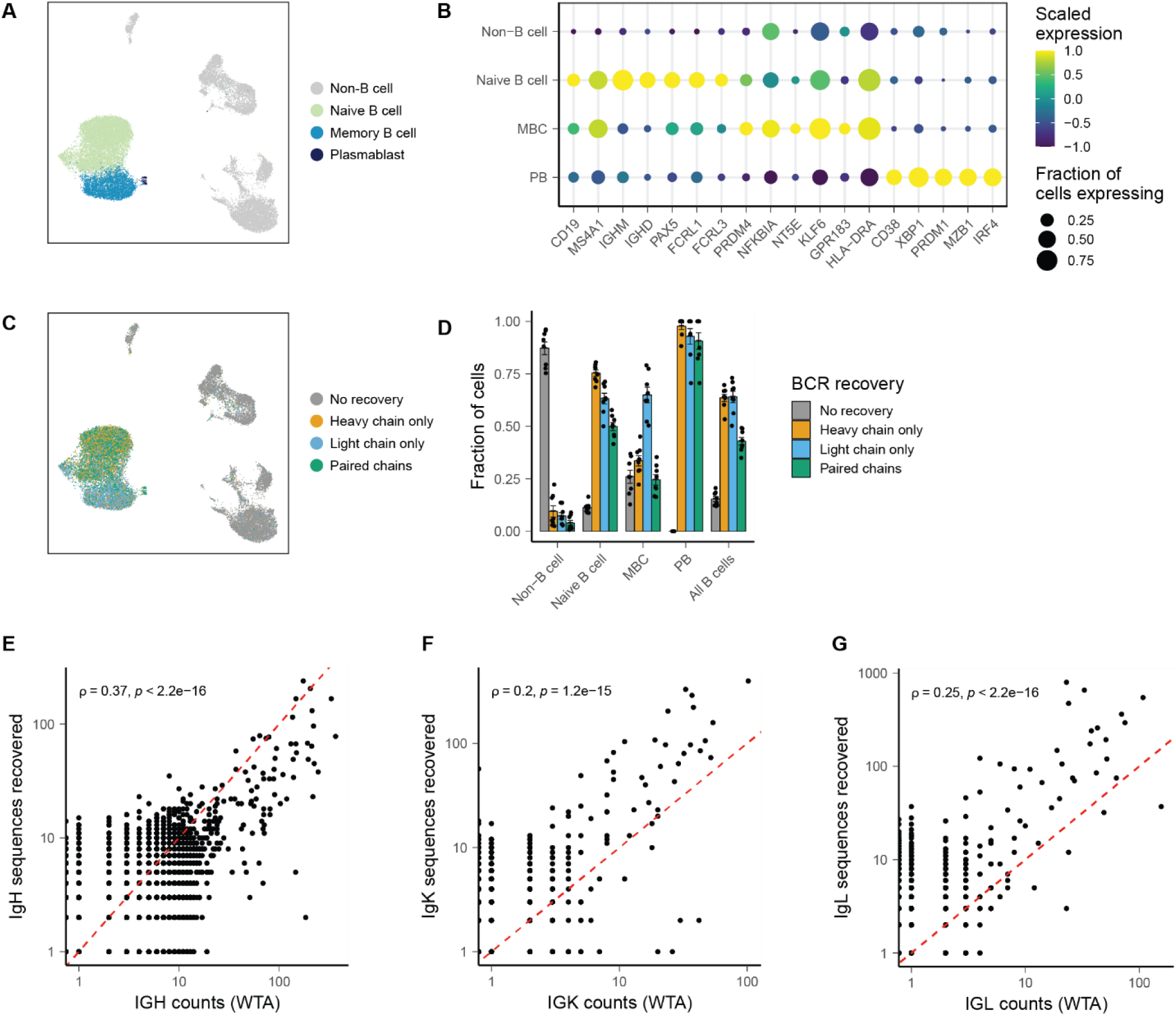
Recovery of full-length, paired BCR sequence from single-cell libraries. A) UMAP of cell phenotypes present in B-cell enriched human PBMC. B) Dot plot showing scaled expression and fraction of cells expressing B cell markers in each phenotype. C) UMAP of BCR recovery from single cells. D) Fraction of cells with no recovery, recovery of heavy chain, recovery of light chain, and paired recovery. Error bars are mean +/- standard error of the mean. E-G) Correlation between number of counts mapping to the IGH/IGK/IGL locus and the number of functional heavy chain or light chain molecules recovered. Spearman’s correlation coefficient and the associated p-value are shown.

We then recovered BCR sequences using our method (Figure 2C). In sum, we recovered functional, full-length heavy chain sequences from 67.8% of B cells, light chain sequences from 57.0% of B cells, and paired heavy and light chain sequences from 39.9% of B cells (Figure 2D). To assess the efficiency of our method, we determined the number of unique molecules mapping to the BCR heavy and light chains in whole transcriptome libraries and compared these counts to the corresponding number of unique molecules recovered from the same cells with our method. We found strong correlations between the number of molecules of heavy chain, kappa, and lambda transcripts in whole transcriptome libraries and the number of unique molecules recovered from our method (Figure 2E-G, Supplemental Figure 1A, 1B). On average, our method recovered full-length variable region for 77.5% as many heavy chain molecules as analyzed in whole-transcriptome sequencing, and 3.0 times (kappa) or 4.97 times (lambda) chain molecules as present in whole-transcriptome sequencing, indicating that our method can recover BCR variable region sequence information from a majority of BCR transcripts present in the starting single-cell WTA product. We attributed the improved ability to recover light chain sequences relative to heavy chain sequences to the overall shorter length of light chain transcripts, which reduces the probability that the variable regions are truncated by the randomly primed second strand synthesis in the library preparation used here and enables an increased cluster density of light chain amplicons on the flow cells used for next-generation sequencing^39, 40^. The recovery of BCRs was dependent the phenotype of the B cell, with elevated levels of recovery from plasmablasts and moderately reduced rates of recovery from MBCs, consistent with differential levels of BCR transcript expression by these cells in our corresponding whole-transcriptome data (Figure 2D, Supplemental Figure 1C, 1D). We also noted that our method appeared to recover BCR sequences from a minority of non-B cells. We further examined the single-cell expression profiles of these cells and found that a majority of these cells contained detectable mapping to BCR heavy and light chains in whole-transcriptome libraries. Indeed, non-B cells with at least one count of heavy chain in WT libraries were 15.9 times more likely than other B cells to have a recovered heavy chain, and non-B cells with at least one count of lambda or kappa chain were 20.3 more times more likely than other B cells to have a recovered light chain (Supplementary Figure 1E, 1F). This finding suggests that the majority of these artifactual sequences were not a product of our method for enrichment, but rather were generated in the preceding steps of single-cell library preparation.

### Recovered BCR sequences demonstrate concordance with single-cell whole-transcriptome libraries

We next examined the agreement between single-cell transcriptomes and our recovered BCR sequences. We first approximated the heavy chain isotypes of single cell transcriptomes using reads mapping to the heavy chain constant region and compared these isotypes to those recovered using our method (Figure 3A). We found that for cells expressing IgM or IgD isotypes in whole-transcriptome sequencing, we recovered a mixture of both IgM and IgD BCRs, while for cells expressing IgG and IgA sequences, we recovered predominantly either IgG or IgA BCRs, respectively (Figure 3B). We also recovered a mixture of IgM and IgD BCRs from cells with a naive B cell phenotype and increasing frequencies of class-switched IgG and IgA BCRs from MBCs and PBs (Figure 3C). These results demonstrate that our method faithfully preserved information related to BCR isotype. We then analyzed the agreement mapping to heavy and light chain V-gene segments present in whole-transcriptome libraries and the V-gene calls recovered by our method. To avoid ambiguities related to differences in the genome and immunoglobulin gene references used in this study, we focused this analysis on a subset of V-genes that existed in both the human genome and IMGT reference used in this study. We found clear trends of concordance between IGHV, IGKV, and IGLV gene usage present in single-cell libraries with the corresponding recovered BCR sequences, suggesting that our method also faithfully recovers information related the variable region of the BCR (Figure 3D-E). Lastly, we computed the somatic mutation frequency of recovered heavy and light chain BCR sequences (Figure 3F). We found clear trends of increasing mutation frequency from naïve to MBC to PB cell phenotypes (Figure 3G-H) and from IgM to IgG and IgA isotypes (Figure 3I). Overall, these results indicate a concordance between the transcriptomes of single cells and the BCR sequences recovered from the same cells with our method.

**Figure 3.**
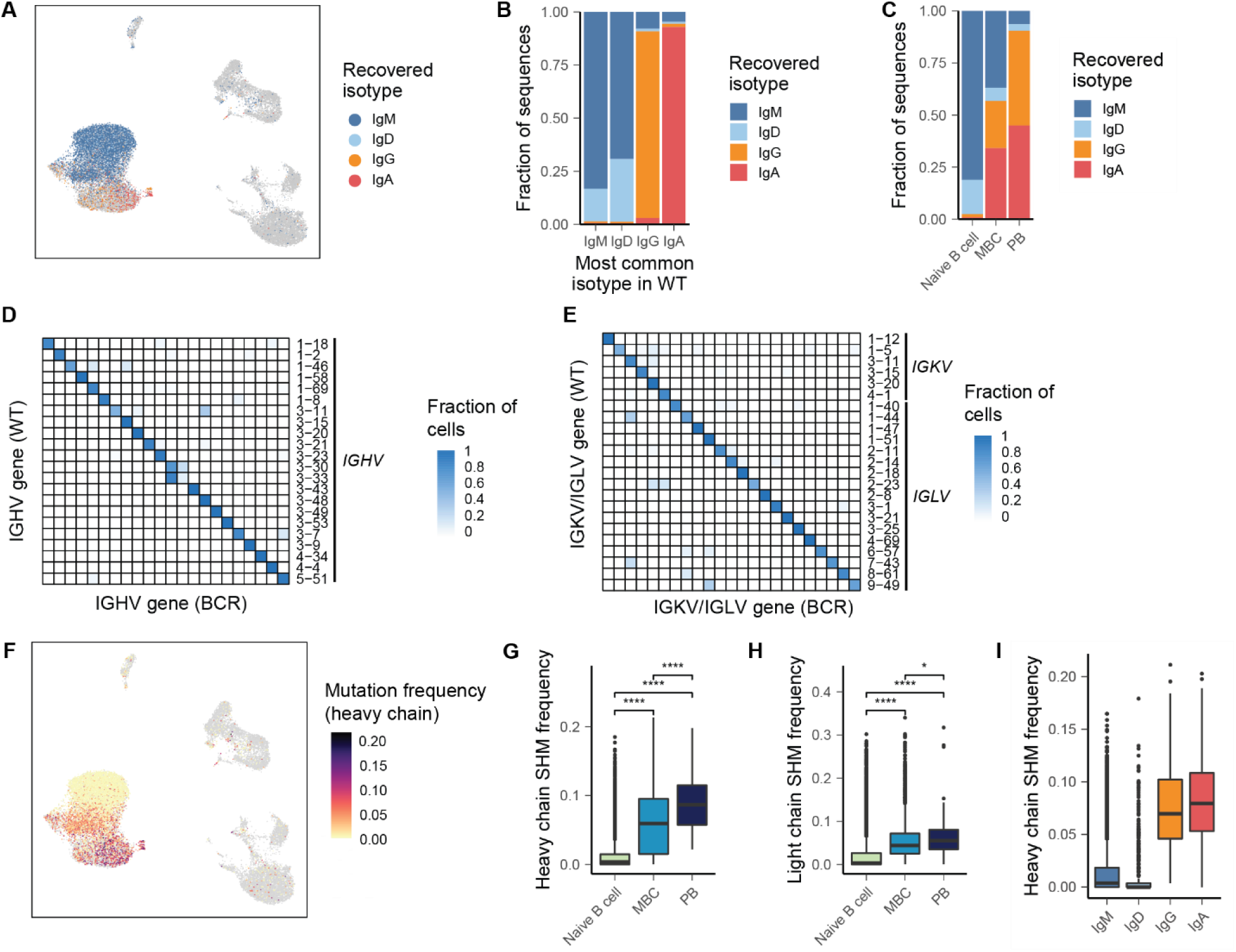
Concordance between single-cell whole transcriptome and BCR libraries. A) Isotypes of BCR heavy chain sequences recovered from single-cell libraries. B) Isotypes of BCR heavy chain sequences recovered from single-cell libraries, grouped by most common isotype present in WT libraries. C) Isotypes of BCR heavy chain sequences recovered from single-cell libraries, grouped by phenotypes assigned in single-cell gene expression data. D-E) Heat map comparing most common heavy chain or light chain V gene segments in WT libraries and V-genes of recovered BCR sequences. F) Heavy chain somatic mutation frequency overlaid onto UMAP. G-H) Somatic mutation frequency of BCR sequences grouped by B cell phenotypes. I) Somatic mutation frequency of BCR sequences grouped by B cell isotypes. P-values are calculated using a two-sided Wilcoxon rank-sum test and are adjusted using Bonferroni correction.

### Recovered BCR sequences demonstrate high levels of accuracy and sensitivity

To further assess the accuracy of the recovered sequences, we also evaluated the ability to recover BCR from a human embryonic kidney (HEK) cell line transiently transfected with separate plasmids encoding the heavy and light chains of a fully human IgG1/κ BCR (U6) (Supplemental Table 1). The next day, we diluted this transfected cell line into B cells isolated from a second sample of PBMC and analyzed the resulting cell suspension with scRNA-seq. In addition to HEK cells, we identified clusters of non-B cells, naive B cells, MBCs, and PBs in the resulting transcriptional data (Figure 4A-B). To verify that the HEK cells expressed BCR transcript, we analyzed the number of heavy and light chain transcripts in WT libraries recovered from each phenotype. Many HEK cells expressed no heavy or light chain transcript in WT libraries, likely owing to poor transfection efficiency under the conditions used or the poor viability of transfected HEK cells relative to non-transfected HEK cells. Nonetheless, we found that about half of HEK cells expressed BCR heavy chain molecules and about 25% of HEK cells expressed BCR light chain transcript (Supplemental Figure 4A-B), indicating the successful transfection of these cells with the U6 BCR.

**Figure 4.**
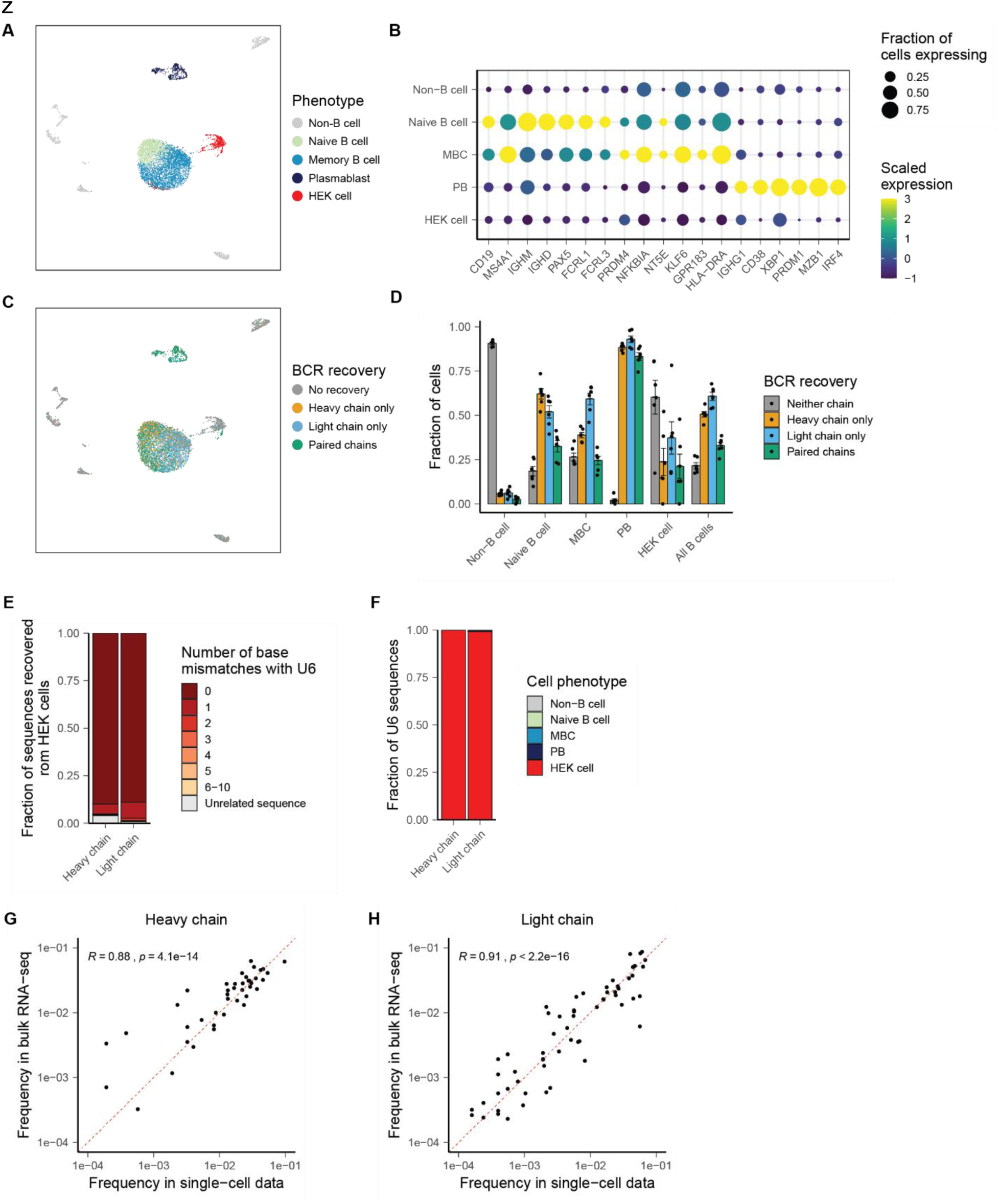
Accuracy and sensitivity of approach. A) UMAP of transcriptional phenotypes present in B cell-enriched PBMC with spike-in of U6-transfected HEK cell line. B) Dot plot showing scaled expression and fraction of cells expressing B cell marker genes. C) UMAP of BCR recovery. D) Rates of BCR recover from each transcriptional phenotype. E) Relationship of heavy and light chain sequences recovered from HEK cells to U6 BCR. F) Cell phenotypes with recovery of U6 BCR. G, H) Correlation between *IGHV* (G) and *IGKV/IGLV* (H) frequency in recovered BCR from single cells and BCR from 5′-RACE bulk sequencing.

We then recovered BCR sequences from these libraries and achieved similar levels of recovery as in our PBMC experiment (Figure 4C-D). We then analyzed BCR sequences originating from HEK cells. Remarkably, 95.2% of heavy chain sequences and 97.4% of light chain sequences recovered from HEK cells exhibited at most one nucleotide mismatch from the heavy or light chain of the U6 BCR (Figure 4C). The remaining sequences recovered from HEK cells either exhibited greater number of nucleotide mismatches with the U6 BCR or were apparently unrelated to the U6 BCR. We hypothesize that these unrelated sequences result either from pre-amplification artifacts (i.e. ambient RNA contamination)^41^ or the formation of chimeric PCR products during amplification^42^. As observed primary with B cells, we found that the probability of BCR recovery from HEK cells was related to the number of molecules enumerated in whole-transcriptome product, suggesting that the inability to recover BCR from many individual HEK cells results from low levels transfection of those same cells or the absence of transfection altogether (Supplemental Figure 2C-D). We also examined the recovery of U6 BCR sequences from all cells and found that greater than 99% of both U6 heavy and light chain sequences were recovered from HEK cells, further confirming that our method is capable of attributing BCR sequences to the correct cells (Figure 4F). These findings demonstrate that our approach possesses high levels of accuracy and sensitivity, allowing the recovery of matching of BCR sequences to single cells with high confidence. In addition, they demonstrate that our approach is highly reliable even with highly oligoclonal populations of B cells, such as clonally expanded, antigen-selected B cells.

Lastly, to assess to what extent our method may exhibit bias in its ability to recover BCRs, we performed bulk VDJ amplicon sequencing of both heavy and light chain of B cells isolated from the same donor, using an approach based on 5′-reversible amplification of cDNA ends (5′-RACE) to avoid introducing any bias that may be associated with amplification with V-region primer sets^43, 44^. We found strong correlations between the frequency of heavy and light chain V-region segments in BCRs recovered from single-cell libraries, confirming that the use of V-region primer sets in our approach does not introduce substantial bias into the sequences recovered (Figure 4G-H).

### Isolation of antigen-specific B cells from rhesus macaques receiving monovalent glycoconjugate pneumococcal vaccines

We next applied our approach to resolve the clonotypic and phenotypic characteristics of antigen-specific B cells elicited by glycoconjugate pneumococcal vaccines. We focused these studies on responses generated against the capsular polysaccharide of *S. pneumoniae* serotype 3 (ST3). We prepared a fluorescently-labelled, biotinylated ST3 polysaccharide to label and isolate ST3-specific B cells. We analyzed PBMC samples from five infant rhesus macaques that were obtained one week after receiving the third dose of a monovalent ST3 glycoconjugate vaccine. We selected these samples from five monkeys, one of which exhibited a strong response to vaccination, as assessed by ST3-specific IgG and opsonophagocytic assay (OPA) titers^45^, two that exhibited intermediate responses, and two of which exhibited low responses. As a control, we also analyzed samples obtained from three adult monkeys that had never received pneumococcal vaccines.

From each monkey, we used fluorescence associated cell sorting (FACS) to isolate ST3-reactive B cells as well as non-ST3-reactive B cells for analysis with scRNA-seq. We found an increase in the frequency of IgG+ ST3-reactive B cells in vaccinated monkeys relative to the control group, indicating that the majority of IgG+ ST3-reactive B cells are elicited by vaccination (Figure 5A-B, Supplemental Figure 3). The small number of ST3-reactive B cells analyzed in unvaccinated monkeys may represent a degree of preexisting humoral immunity acquired through exposure to similar antigens, such as those present in the gut flora^46^, though it remains possible that a fraction of these B cells binds to the antigen construct through non-specific means. We also observed modest correlations between the frequency of IgG+ ST3-reactive B cells and ST3-specfic IgG and OPA titers, providing support for our strategy to isolate ST3-reactive cells and suggesting that stronger responses to the ST3 antigen are linked to higher frequencies of class-switched, ST3-reactive B cells (Figure 5C-D).

**Figure 5.**
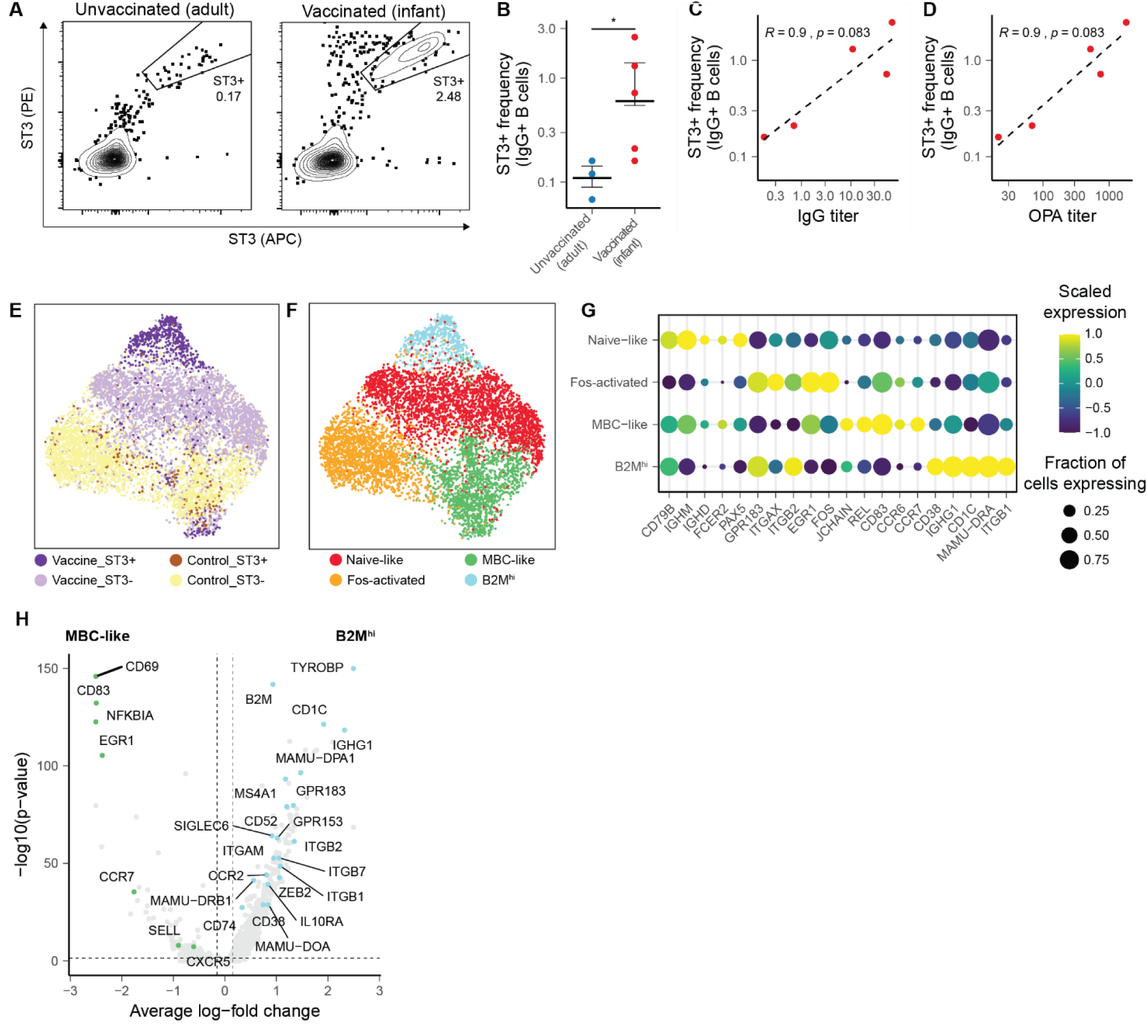
Transcriptional features of ST3-reactive and non-ST3-reactive B cells elicited by vaccination. A) Representative staining of ST3-reactive B cells from unvaccinated and vaccinated samples, gated on IgG+ B cells (Live CD3^-^ CD19^+^ CD20^+^ IgG^+^). B) Frequency of ST3-reactive IgG+ B cells in vaccinated and unvaccinated monkeys. C, D) Correlation between ST3-reactive IgG+ B cells and ST3-specific IgG (C) and OPA (D) titers. E, F) UMAP of single-cell transcriptomes colored by sort fraction (E) and cell phenotype (F). Dot plot showing scaled expression and percent of cells expressing B cell marker genes in each transcriptional phenotype. H) Volcano plot of differentially expressed transcripts between B2M^hi^ and MBC-like cells. P-values are calculated using a two-sided Wilcoxon rank-sum test and are adjusted using Bonferroni correction.

Across all 8 animals, we recovered high quality single-cell transcriptomes for 6,819 cells, including 947 ST3-reactive cells (Figure 5E). In the resulting transcriptional data, we annotated four clusters of B cells, including: naïve-like B cells, Fos-activated B cells, which upregulated transcripts associated with the transcription factor AP-1 (*FOS, FOSB, JUNB*), memory-like B cells (MBC-like cells), as well as an additional population of memory B cells (B2M^hi^) which exhibited a downregulation of markers associated with lymph homing (*SELL, CCR7*, *CXCR5*), elevated levels of BCR inhibitory molecules (*SIGLEC6*, *SIGLEC10*), and an upregulation of molecules associated with tissue homing (*ITGAX, ITGB2, ITGB7*)^25, 47^. Compared to B2M^hi^ B cells, MBC-like cells also upregulated transcripts associated with immunological memory (*BCL2A1*), as well as *NF*-kB signaling (*NFKBIA, NFKBID, REL, RELB*, while B2M^hi^ cells upregulated transcripts associated with antigen presentation (*CD74, MAMU-DRA*, *MAMU-DPA1, MAMU-DOA, MAMU-DRB1*) and with B cell activation (*IGHG1, GPR183, CD1C, MS4A1*) (Figure 5H, Supplemental Table 2)^48, 49^.

### Select features of the BCR repertoire are associated with specificity for ST3

To assess the repertoire of ST3-reactive B cells, we next recovered full-length variable region BCR sequences (Supplemental Figure 4A-B). The degree of BCR recovery achieved was slightly higher among ST3-reactive cells but did not differ substantially among transcriptional phenotypes (Supplemental Figure 4C). Consistent with our cluster annotations, we observed greater levels of somatic hypermutation frequency among both MBC-like cells and B2M^hi^ cells than naïve cells, as well as increased class switching to IgG among B2M^hi^ cells (Figure 6A-D, Supplemental Figure 4D). We observed an increase in the fraction of IgG sequences among ST3-reactive cells in four of the five vaccinated monkeys analyzed, which could indicate enhanced class-switching promoted by vaccination with the monovalent glycoconjugate vaccine used here (Supplemental Figure 4E).

**Figure 6.**
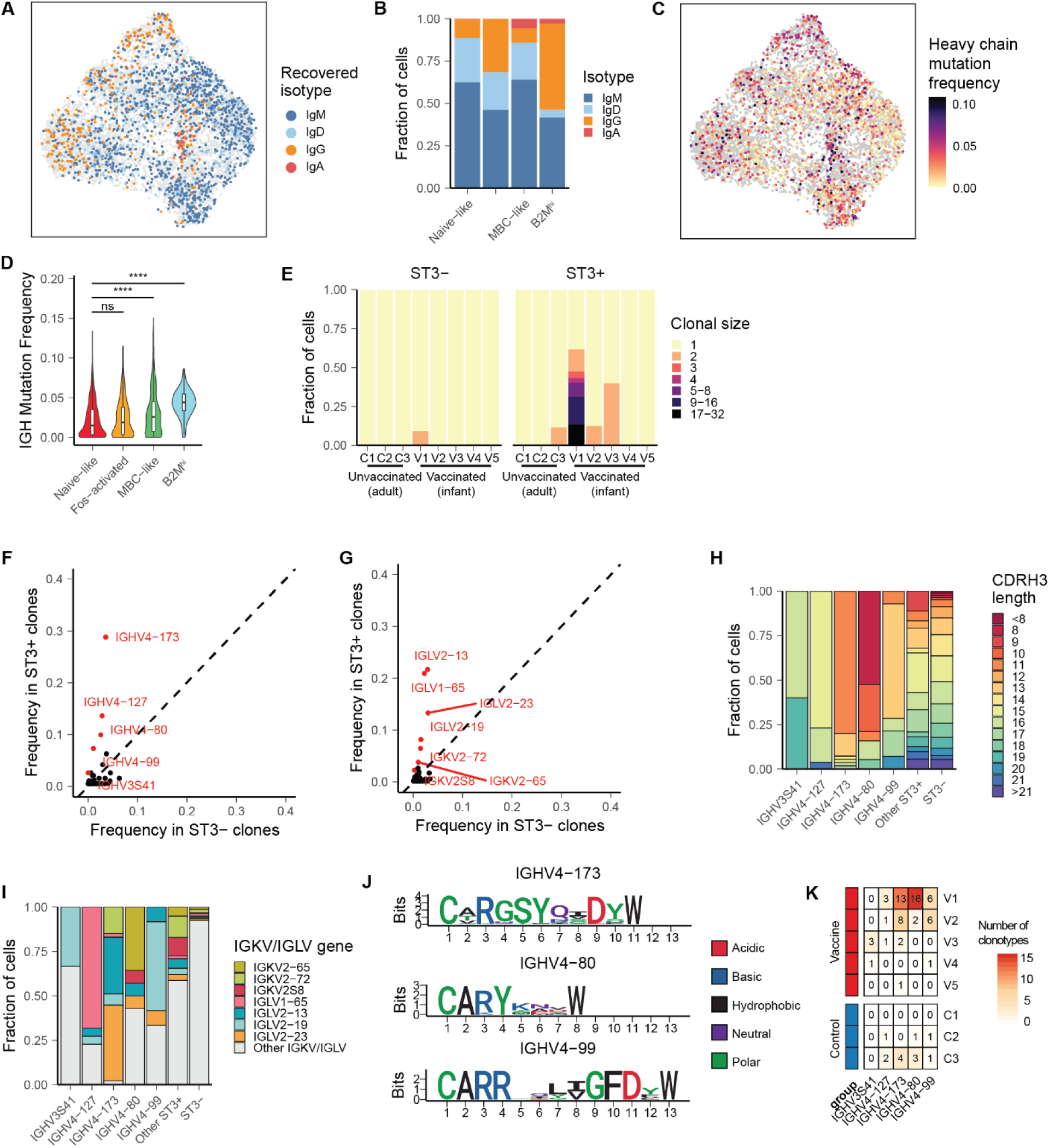
Clonotypic features of ST3-reactive B cells. A) UMAP colored by isotype of recovered BCR. B) Isotypes recovered from B cell phenotypes. C) UMAP colored by frequency of somatic mutation. D) Frequency of somatic mutation in heavy chain sequences recovered from each phenotype. P-values are calculated with a two-sided Wilcoxon rank-sum test and are adjusted with Bonferroni correction. E) Clonal sizes of ST3-reactive and ST3-B cells. F, G) Frequency of IGHV (F) and IGKV/IGLV (G) genes among ST3+ cells from vaccinated monkeys and all ST3-cells. Genes highlighted in red are statistically significantly enriched in the ST3+ fraction (p < 0.001, p-values calculated using a two-sided chi-squared test and are adjusted using Bonferroni correction). H) CDRH3 junction lengths of ST3-reactive cells using *IGHV* genes statistically associated with ST3-reactive cells. I) IGKV/IGLV gene pairings lengths of ST3-reactive cells using *IGHV* genes statistically associated with ST3-reactive cells. J) Logo plots of CDRH3 junctions of ST3-reactive clonotypes using IGHV4-173, IGHV4-80, and IGHV4-99. K) Number of clonotypes with each V-gene recovered from each vaccinated and unvaccinated monkey.

Next, we defined clonal lineages using the recovered heavy chain sequences. We found a substantial increase in the frequency and size of expanded clonal lineages recovered from ST3-reactive cells in three of the five vaccinated monkeys, consistent with these cells having recently undergone expansion in response to antigen encounter (Figure 6E). Expanded clonotypes in other cells were likely not detected in the other two vaccinated monkeys due to the low number of ST3-reactive cells from these monkeys for which BCR information was recovered (Supplemental Figure 4E-F). All expanded clonal lineages were detected exclusively in either the ST3+ or ST3-fractions, confirming that reactivity to the ST3 antigen is shared by cells in a clonotype and supporting that our cell isolation strategy successfully isolated ST3-specific B cells from non-ST3-specific B cells (Supplemental Figure 4G). Interestingly, two expanded ST3+ clonotypes were detected in a single monkey from the unvaccinated control group (Supplemental Table 3). These clonotypes expressed IgM isotypes but had BCRs that were not in germline configuration, suggesting that they had undergone mutation in response to prior exposure to a similar antigen.

We next aimed to define BCR motifs associated with specificity for the ST3 antigen. First, we compared the frequency of V gene segments utilized by heavy and light chain sequences between ST3-reactive cells from the vaccine group and non-ST3-reactive cells. We determined that five IGHV genes and seven IGKV or IGLV genes were statistically enriched (Bonferroni-adjusted p-value < 0.001) in frequency among ST3+ cells, suggesting that these gene segments encode motifs that promote specificity to the ST3 antigen (Figure 6F-G). Together, this set of V genes comprised 62.3% of all ST3-reactive heavy chain sequences and 75.3% of ST3-reactive light chain sequences recovered. To further define motifs associated with ST3 reactivity, we grouped ST3-reactive sequences by these IGHV genes and analyzed the lengths of their CDRH3 junctions and the usage of IGKV/IGLV genes by their paired light chains (Figure 6H-I). We found a clear preference for CDRH3 length and IGKV/IGLV genes by ST3-reactive clonotypes utilizing each of these IGHV genes, suggesting that these combinations of BCR features are associated with reactivity towards ST3. For the three IGHV genes for which we recovered the greatest number of unique clonotypes, we further analyzed CDRH3 junctions and identified further sequence characteristics associated with reactivity to ST3, including: “GSY” at codons 4-6 of the CDRH3 junction of ST3-reactive clonotypes using IGHV4-173, “Y” at codon 4 of clonotypes using IGHV4-80, and “L(V/I)G” at codons 9-11 of clonotypes using IGHV4-99 (Figure 6J).

### Rhesus macaques exhibit a convergent repertoire in response to vaccination

We next assessed to what extent these sequences motifs were utilized by each of the vaccinated monkeys. Remarkably, all five IGHV genes identified here as associated with ST3-reactivity were utilized by clonotypes present in 2 or more monkeys in the vaccine group, and all five vaccinated monkeys possessed clonotypes with at least one of these IGHV genes, indicating a convergence for a defined set of ST3-reactive sequences in monkeys receiving this pneumococcal vaccine formulation (Figure 6K). Interestingly, we also identified clonotypes that utilized four of these five IGHV genes among ST3+ cells from two of the three unvaccinated monkeys. We further examined these potentially ST3-reactive sequences from unvaccinated monkeys and found that, while IGHV4-127, IGHV4-80, and IGHV4-99 from these unvaccinated monkeys exhibited preferences for distinct CDRH3 junction lengths and IGKV/IGLV gene pairings than corresponding ST3-reactive sequences from vaccinated monkeys, a subset of IGHV4-173 ST3-reactive sequences from unvaccinated monkeys exhibited a similar preference for CDRH3 junctions of 11 or 12 amino acids as well as pairing with IGKV2-65 or IGLV2-23, demonstrating a high level of similarity with ST3-reactive clonotypes recovered from vaccinated monkeys (Supplemental Figure 5A-B). Consistent with this observation, a recent study of patients receiving the 23-valent Pneumovax polysaccharide pneumococcal vaccine demonstrated that the vaccine resulted in the expansion of pre-existing capsular polysaccharide-specific plasma cells that shared reactivity with antigens present in the gut flora^46^. Overall, these findings demonstrate a convergent response to vaccination among the infant monkeys studied, in which each vaccinated monkey possesses BCR clonotypes selected from a common set of features. In addition, our data suggests that unvaccinated adult monkeys may possess B cells that share BCR features with clonotypes elicited by the monovalent ST3 glycoconjugate vaccine studied here.

## Discussion

Here, we have described a method for the reliable recovery of full-length, paired BCR sequences from 3′-barcoded scRNA-seq libraries. We demonstrated that this approach allows for the recovery of accurate sequences with a strong concordance to the single-cell transcriptomes of the same cells. The reagents and platforms necessary to implement our method are inexpensive and widely available. Importantly, our method can be used to sequence BCR variable regions for both new and preexisting 3′-barcoded single-cell libraries.

Using this method, we analyzed B cell responses elicited against the ST3 antigen by glycoconjugate vaccines. Using our approach, we reveal clonotypic features associated with specificity against the ST3 antigen, and we show that these features emerge in clonotypes recovered from distinct vaccinated monkeys, demonstrating a convergence towards a common set of antigen-specific motifs. We hypothesize that the convergence of these heavy and light chain paratopes results in the recognition of common, immunodominant epitopes present on the ST3 antigen, similar to what has been observed in other contexts^18, 50–54^. Interestingly, the simple structure of the ST3 antigen, compared to the polysaccharide capsules of other pneumococcal serotypes, may influence the degree of repertoire convergence by exposing a limited number of epitopes^55, 56^. Further studies can assess the importance of these clonotypic and transcriptional features in providing long-lasting immune protection against ST3 and can compare these results to those generated with vaccines against other pneumococcal polysaccharides.

Our method has at least two practical limitations. First, it does not recover paired, full-length BCR for every B cell present in a pooled scRNA-seq product. This limitation likely derives from limitations in the efficiency of RNA capture inherent to the scRNA-seq platforms utilized here. Indeed, it is well-established that scRNA-seq libraries contain only a fraction of the total mRNA content of a single cell^35, 57, 58^, and here we have consistently observed correlations between levels of BCR transcript expression and the probability of BCR recovery from the same cell. Thus, future advances in the molecular biology used to prepare scRNA-seq libraries could enhance the sensitivity of our approach. In addition, our method requires the use of species-specific elements, including the multiplexed V-region primer sets used for primer extension. The design of these primers requires the accurate annotation of V genes for a given organism. While primer sets for many commonly studied model organisms are available^59–63^, this requirement may prove a barrier to studies of less well-characterized organisms. Despite this limitation, the characterization of V genes in many species continues to improve, and we expect the availability of suitable primer sets to continue to expand in the future^61, 64, 65^.

In summary, we have described an approach for the reliable recovery of full-length, paired heavy and light chain BCR sequences from 3′-barcoded scRNA-seq libraries. The method exhibits high sensitivity and recovers sequences that exhibit high levels of accuracy and concordance with the transcriptional profiles of the same cells, and the sensitivity of the method is limited predominantly by the efficiency of mRNA capture and level of BCR expression. By applying the method to antigen-specific B cells elicited by pneumococcal vaccination, we demonstrate a convergence of BCR features among distinct monkeys. We anticipate that this method will be especially useful in analyzing the relationships between phenotypic and clonotypic features of antigen-specific B cell populations in vaccination or disease.

## Materials and Methods

### Ethics statement

Immunizations for the non-human primate (NHP) study were performed at the University of Louisiana at Lafayette-New Iberia Research Center (NIRC), which is accredited by the Association for Assessment and Accreditation of Laboratory Animal Care (AAALAC, Animal Assurance #: 000452). The work was in accordance with USDA Animal Welfare Act and Regulations and the NIH Guidelines for Research Involving Recombinant DNA Molecules, and Biosafety in Microbiological and Biomedical Laboratories. All procedures performed on these animals were in accordance with regulations and established guidelines and were reviewed and approved by an Institutional Animal Care and Use Committee or through an ethical review process.

### Animals and immunization

Infant rhesus macaques were housed with their moms for the duration of the study at the New Iberia Research Center (NIRC), University of Louisiana. infant rhesus macaque experimental protocol was approved by IACUC at both Pfizer Inc and NIRC. Age (3-6 months old at the start of the study) and sex matched infant rhesus macaques were randomly divided into groups for vaccination. Infants were vaccinated under sedation with 2.2 µg of pneumococcal serotype 3 conjugate vaccine (ST3) formulated with 125 µg of AlPO_4_ at 0.25 ml volume of final formulation intramuscularly (IM) into a single limb on day 0. Pre-bleeds to assess baseline ST3-specific sera titers were collected 1 week (wk= -1) before primary vaccination (D0). Post vaccination blood for sera were collected at 4-weeks post-dose 1 (PD1).

Control serum and PBMC samples were collected from adult Rhesus Macaques at NIRC from naïve animals which have not been previously vaccinated with pneumococcal vaccines.

### Determination of ST3-specific IgG and OPA titers

Streptococcus pneumoniae direct Luminex immunoassays (dLIAs) were used to quantify IgG antibody concentrations in serum for ST3 polysaccharide. This was done using Luminex Magplex technology (Luminex Corporation, Austin, TX). Briefly, serum samples, reference standard serum and QCS prepped by Hamilton Robotic units (Hamilton Company, Reno, NV). PnPS-coupled magnetic microspheres were added to each well of the assay plates and incubated. After 90 minutes non-bound components were washed off and the secondary antibody (R-Phycoerythrin-conjugated goat anti-human IgG) was added to each well. After 90 minutes the assay plate was washed again and 100ul of LXA-20 was added to each well. After 2-hours of incubation, the plates were then loaded into Bio-plex 200 and median fluorescent intensities (MFI) were read. These MFIs were used to calculate antigen specific IgG concentrations (µg/ml) against established IgG concentrations of the reference standard by statistical analysis software (SAS).

Microcolony OPA is based on the antibody-mediated complement- and phagocyte-dependent OPA assays as described previously. Here, OPA assays were developed for S. pneumoniae serotype 3. Briefly, heat-inactivated sera were serially diluted. Target bacteria were added to assay plates and incubated. Baby rabbit complement and differentiated HL-60 cells were added to each well at an approximate effector to target ratio. After incubations, resulting colonies were stained with Coomassie Brilliant Blue stain. Colonies were imaged and enumerated on a CTL ImmunoSpot Analyzer®. The interpolated OPA antibody titer is the reciprocal of the dilution that yields a 50% reduction in the number of bacterial colonies when compared to control wells that did not contain serum.

### Preparation of biotinylated ST3 antigen

Pneumo serotype 3 polysaccharide was mixed with imidazole (3x, w/w) and pH was adjusted to 3.5 with 1M-HCl, then frozen and lyophilized.

After lyophilization, the lyophilized polysaccharide was reconstituted with anhydrous DMSO to make 4 mg/mL saccharide concentration. The reaction mixture was then warmed to 35 C, and subsequently CDI (0.1 MEq) was added. The reaction mixture was stirred at 35 C for 3 hrs. After the reaction mixture was cooled to 23 C, WFI (2% v/v) was added to quench free CDI and stirred further for 30 min at 23 C. Biotin hydrazide (1 MEq) was added. The reaction mixture was stirred at 23 C. After 20 hrs reaction, the reaction mixture was diluted to chilled (at 5 C) PBS buffer (5X, v/v). The diluted reaction mixture was then purified by UF/DF using 10K MWCO PES membrane (Millipore Pellicon 2 Mini) against PBS buffer (30X, v/v) and filtered through 0.22 um and analyzed.

### PBMC isolation

Venous blood samples were collected in heparinized tubes. Each sample was diluted 1:1 with 0.9% NaCl and applied to Accuspin tubes (Sigma-Aldrich, Burlington, MA) filled with a Ficoll– Paque solution (ThermoFisher Scientific, Waltham, MA) before centrifugation. PBMCs were recovered from the interface, washed once in 1X HBSS Cellgro (ThermoFisher), resuspended in Recovery Cell Culture Freezing Medium (Gibco) at a concentration of 5×10^6^ cells/ml and frozen in liquid nitrogen for future use.

### ST3-PE and ST3-APC coupling

Biotinylated pneumococcal serotype 3 polysaccharide with a 5% biotin load was coupled with high concentration streptavidin-phycoerythrin (SA-PE) and streptavidin-allophycocyanin (SA-APC, both from BioLegend, San Diego, CA) in separate reactions. Biotinylated ST3 and the fluorophore-conjugated streptavidin reagents were prepared with PBS at 4X working solutions of 20 µg/mL and 200 µg/mL, respectively. Equal volumes of ST3 and SA-PE; and, separately, equal volumes of ST3 and SA-APC working solutions were combined, thoroughly mixed and incubated at 4 °C protected from light for a minimum incubation period of 60 minutes with thorough mixing at 30 minutes and directly prior to use. Coupled ST3-PE and ST3-APC solutions were maintained at 4 °C, protected from light until use.

### Magnetic isolation of B cells from human PBMC

Whole human blood was obtained from Research Blood Components, LLC (Watertown, MA), and PBMC were isolated by means of density gradient centrifugation and cryopreserved. Upon use, PBMC were thawed into Aim-V medium (ThermoFisher). B cells were isolated using an EasySep Human Pan-B Cell Enrichment kit (Stemcell Technologies, Vancouver, Canada).

### Plasmid generation

Monoclonal antibody U6 heavy and light chain vectors were created from an affinity-selected Ara h 2-reactive B cell from a patient undergoing peanut oral immunotherapy (NCT01324401), as previously described^17^. Briefly, peripheral blood cells were isolated by Ficoll gradient (GE Healthcare) and stained with an Ara h 2-Alexa Fluor 488 multimer and fluorescent antibodies, CD3–APC (eBioscience), CD14-APC (eBioscience), CD16-APC (eBioscience), CD19–APC-Cy7 (BD Biosciences), CD27–PE (BD PharMingen), CD38–BV421 (BD Biosciences), and IgM–PE-Cy5 (BD PharMingen), for single-cell sorting on an FACS Aria II sorter (BD Biosciences) into a 96 well plate. Heavy and light chains underwent nested amplification using previously published primers sets^17^ followed by Sanger sequencing (Genewiz). Productive sequences were cloned into a heavy chain IgG1 vector and a light chain kappa vector using restriction enzymes AgeI, BsiWI, Sall, and XhoI (New England Biolabs) for transformation with competent Escherichia coli NEB5a bacteria (New England Biolabs) and colony selection. Plasmid DNA from overnight liquid cultures (LB with 100 mg/ mL ampicillin) was isolated with QIAprep Spin Miniprep Kit (Qiagen).

### HEK cell culture and transfection

Plasmid DNA (25 ng) was transfected into HEK293 T cells (ATCC, CRL-3216) by using GenJet In Vitro DNA Transfection Reagent (SignaGen, Rockville, Md) and cultured overnight at 37C culture at 378C with 5% CO2 in serum free HL-1 media (Lonza, Walkersville, Md) supplemented with Pen-Strep and 8 mmol/L Glutamax (Gibco, Carlsbad, Calif).

### Flow cytometry and cell sorting

Cryopreserved PBMC were thawed in AIM-V media and washed twice with PBS. Tetramer reagent was prepared by combining equal volumes of streptavidin-PE or streptavidin-APC solutions (both from BD) at 200 ug/mL with polysaccharide-biotin solution at 20 ug/uL. Cells were then stained with eBioscience Fixable Viablity Dye eFlour 780 (ThermoFisher) and Fc block (BD) on ice for 30 minutes. Cells were then washed twice with PBS and then stained with 25 uL of each ST3-PE tetramer and ST3-APC tetramer on ice for 30 minutes. Cells were then washed three times in FBS staining buffer (BD) and were then stained with BV605-conjugated anti-IgM (MHM-88), BV786-conjugated anti-CD20 (both from Biolegend), DyLight 405-congjuatated anti-CD19 (CB19) (R&D systems), V500-conjugated CD3 (SP34-2), PE-Cy5-conjugated anti-IgG (G18-145) (both from BD), and Total-seq A anti-human hashtag antibody (Biolegend) on ice for 30 minutes. Live+CD3-CD19+CD20+ST3+ and Live+CD3-CD19+CD20+ST3-cells were sorted on a BD Aria 3 sorter, and data was analyzed using Flowjo v10.

### Single-cell RNA-sequencing

Single-cell suspensions from esophageal biopsies, duodenal biopsies, and sorted subsets of peripheral blood CD4+ memory T cells were processed for single-cell RNA sequencing using the Seq-Well platform with second strand chemistry, as previously described^36, 40^. Libraries were barcoded and amplified using the Nextera XT kit (Illumina, San Diego, CA) and were sequenced on a Novaseq 6000 (Illumina).

### Analysis of single-cell whole-transcriptome data

Raw read processing of scRNA-seq reads was performed as previously described^35^. Briefly, reads were aligned to the hg38 reference genome or Mmul_10 reference genome and collapsed by cell barcode and unique molecular identifier (UMI). Whole transcriptome data was further analyzed in Seurat^66^. First, cells with less than 300 unique genes detected and genes detected in fewer than 5 cells were filtered out. Variable features were then determined using the FindVariableFeatures function, and the ScaleData function was then used to regress out the number of RNA features in each cell. The number of principal components used for visualization was determined by examination of the elbow plot, and two-dimensional embeddings were generated using uniform manifold approximation and projection (UMAP). Clusters were determined using Louvain clustering, as implemented in the FindClusters function in Seurat. For analysis of ST3-reactive B cell data, clusters of cells that exhibited high expression of mitochondrial genes or markers associated with other cell phenotypes (i.e. monocytes) were moved and the data was reprocessed. Differential gene expression analysis was performed for each cluster and between indicated cell populations using the FindMarkers function.

### Enrichment of BCR transcripts

Enrichment of BCR transcripts from whole transcriptome amplification products generated with Seq-Well was performed using xGen Lockdown reagents (IDT). Biotinylated probes for IGHM, IGHD, IGHG, IGHD, IGHE, IGKC, and IGLC were synthesized by IDT and were used at a concentration of 1.5 uM (each oligo) (Supplemental Table 4). 3.5 uL of WTA product was combined with 8.5 uL 2x hybriziation buffer, 2.7 uL hybridization buffer enhancer, 0.8 uL Seq-Well WTA primer (40 uM), and 0.5 uL human cot-1 DNA (IDT) were combined and incubated at 95C for 10 minutes. Then, 1 uL of hybridization probe mix was added. The mixture was incubated at 65C for one hour, and was then processed according to the remainder of the xGen lockdown protocol. 50 uL of streptavidin M270 Dynabeads (ThermoFisher) were used for each sample. At the end of this protocol, products were eluted in 20 uL.

The enriched product was then amplified with PCR. Five PCR reactions for each enriched sample were performed with the following composition per reaction: 12.5 uL of 2x Kapa Hifi Hotstart Readymix (Roche), 8.5 uL water, 2.0 uL Seq-Well WTA primer (10 uM), and 2.0 uL of BCR-enriched product. The following PCR cycling conditions were used: 1 cycle of 95C, for 3 minutes; 25 cycles of 98C for 40s, 67C for 20s, 72C for 1 min; 1 cycle of 72C for 5 min. The reactions were then pooled and purified using a homemade SPRI reagent at a ratio of 0.80x^67^.

### Construction and sequencing of BCR sequencing libraries

We designed and constructed IGHV-Nextera, IGKV-Nextera, and IGLV-Nextera primers for both human and rhesus macaque. For rhesus macaque, we used modified sets primers adapted from Rosenfeld et al^59^. For human and rhesus macaque, heavy chain primers were based on the BIOMED-2 primer system^60^. For human light chain, primers were adapted from Reddy et al^20^. (U6 spike-in experiment) or Jiang et al. (PBMC data in Figures 2 and 3)^68^ (Supplemental Table 4). Reaction mixtures for heavy and light chain were composed of: 12.5 uL 2x Kapa Hifi Hotstart PCR Readymix, 6.0 uL water, 4.08 uL primer mix, and 4.0 uL of enriched product. Primer extension was performed with the following thermal program: 98C, 5 minutes, 55C, 30s, 72C, 2 min. The products were then cleaned with SPRI at a ratio of 0.8x (light chain) or 0.65x (heavy chain) and were eluted into 11 uL of water.

Four reactions of library PCR were performed per sample. Reactions were composed of: 0.5 uL P5-Seq-Well primer (10 uM), 0.5 uL P7-Nextera primer (10 uM), 12.5 uL 2x Kapa Hifi Hotstart Readymix, 9 uL of water, 2.5 uL of primer extension product. Amplification used the following cycling conditions: 1 cycle, 95C, 2 min; 14-20 cycles of 95C, 30 s, 60C, 30s, 72C, 1.5 min; 1 cycle of 72C, 5 min. Reactions were pooled and purified using SPRI at a ratio of 0.65x (heavy chain) or 0.8x (light chain). Final products were assessed using an Agilent Tapestation. Light chain libraries usually show a clean peak around 1000bp, and heavy chain libraries usually show a clean peak around 1600bp. Libraries were sequenced on an Illumina Miseq using 600 cycle kits or on an Illumina Novaseq 600 using 500 cycle kits. First, the Seq-Well R1 primer was used to sequence the cell barcode and UMI (20bp on Miseq, 26bp on Novaseq). Then, custom sequencing primer specific for the BCR constant region were used to sequence the BCR using the index 1 read (300bp on Miseq, 270 bp on Novaseq) (Supplemental Table 4). The 8 base pair-length i5 index barcode was sequenced using the Seq-Well R2 index primer (Novaseq) or no custom primer (Miseq). Lastly, the Nextera primer was used to sequence the remainder of the BCR with read 2 (300bp on Miseq, 220 bp on Novaseq).

### Processing of raw single-cell BCR sequencing data

Processing of BCR sequence data was performed with pRESTO and Change-O from the Immcantation software suite^69, 70^. First, potential errors in cell barcode and UMI sequence were corrected for errors up to one nucleotide mismatch with a directional UMI collapse, as implemented in UMI-Tools^71^. Data was filtered by Q-score using the FilterSeq.py function to remove any sequences with an average Q score below 25. Then, the MaskPrimers.py function was used to annotate sequences with the correct isotype (R1 index) and V-region (R2) primer, as well as to mask the corresponding primer regions. The PairSeq.py function was used to retain only read pairs that passed both the FilterSeq.py and MaskPrimers.py processing steps. The data was then segregated by cell barcode and UMI, and the BuildConsensus.py function was used to call consensus sequences for read 1 index and read 2 of each barcode and UMI separately. The AssemblePairs.py function was used to assemble these consensus sequences into a single overlapping sequence. The resulting sequences were then analyzed with IgBlast^72^ using reference sequences provided by IMGT.

### Downstream processing of single-cell BCR data

Molecular consensus sequences were filtered based on the total number of reads (separate for each dataset): >5 reads: rhesus macaque data; >10 reads: PBMC data, heavy chain for HEK experiment; >20 reads: light chain for HEK experiment). Molecular consensus sequences that contained greater than four ambiguous “N” characters were also discarded. BCR sequences were matched to single-cells using the 12 bp single-cell barcodes with a 1 nucleotide Hamming distance error tolerance. To determine a single consensus sequence for each cell, we first considered all sequences attributed to a single cell. If multiple sequences were recovered, we determined a cell-level consensus sequence as follows: first, we performed single-linkage clustering on the recovered IMGT-gapped sequences using Levenstein distance and cut the resulting dendrogram at a height of five to produce clusters of closely related sequences. We further considered sequences that belonged to the largest cluster and had uniform length and determined a gapped consensus sequence. If the resulting consensus retain ambiguous (“N”) characters, we discarded the sequence that was supported by the fewest number of reads and re-attempted to determine a consensus sequence. This process allowed us to combine information from multiple BCR molecules recovered from the same cell into a single cell-level consensus sequence.

### Analysis of B cell clonotypes

Clonotypes were determined using the DefineClones.py function in Change-o^70^, using a Hamming distance calculation and a similarity threshold of 0.10. Germline sequences were then constructed using the CreateGermlines.py function, and frequencies of SHM were determined using the observedMutations function, implemented in Shazam, using the predicted germline sequence with a masked D gene.

### Bulk BCR sequencing

Bulk heavy chain variable region sequencing was performed using 5′-RACE. RNA from magnetically isolated B cells was extracted using the Nucleospin XS RNA kit (Machery-Nagel) and eluted into 12 uL of RNAse-free water and stored at -80C until use. To perform reverse transcription (RT), 6 uL of RNA was combined with 2 uL of primer mix containing 20 µM each of RT primers for IgM, IgG, IgA, IgK, and IgL and 400 U/uL NxGen RNAse inhibitor (LGC Biosearch). This mixture was then heated to 70C for 4 minutes, cooled to 42 degrees for 3 minutes, and cooled to 4C. The RT reaction mixture was prepared on ice and contained (per sample): 4 µL SmartScribe 1^st^ Strand 5X buffer (Takara), 2 µL 100 µM DTT (Takara), 2 µL 10 µM template switching oligo, 2 µL 10 mM dNTPs (New England Biosciences), and 2 µL Smartscribe Reverse Transcriptase (Takara). RNA mixtures were combined with the RT reaction mixture, and RT was performed at 42C for 90 minutes, followed by heating to 70C for 10 minutes and cooling to 4C. PCR1 was performed separately for light and heavy chain; the reaction mixture was then prepared as follows: 5 uL RT product, 5 uL 5X Phusion Buffer (NEB), 10 uL 1M Trehalose (Life Sciences Advanced Technologies), 2.5 uL 10 µM 5′-PCR1 primer, 2.5 uL each PCR1 3′-constant region primer, 1 µL Phusion polymerase (NEB), and remainder water (to 50 µL). PCR1 was performed as: 1 cycle, 98C, 1 minute; 25 cycles: 98C for 20s, 58C for 30s, 72C for 1 minute; 1 cycle, 72C for 1 minute, hold at 4C. PCR2 reaction mixture was performed as follows: 18 µL water, 25 uL 2X Kapa HiFi HotStart Readymix (Roche), 1.5 µL 5′-PCR2 primer (10 µM), 1.5 uL 3′-PCR2 primer (10 µM). PCR2 was performed as follows: 14-18 cycles of: 98C for 40s, 65C for 20s, 72C for 30s; 1 cycle of 72C for 5 minutes, hold at 4C. Library quality was assessed using an Agilent Tapestation D5000 assay, and libraries were sequenced on an Illumina Novaseq SP 500 cycle kit, using 250 x 250 bp reads. All primers were purchased from Integrated DNA Technologies (IDT). Primer sequences used for bulk BCR sequencing are contained in Supplementary Table 5.

### Processing of bulk BCR data

Processing of bulk BCR sequence data was performed with pRESTO and Change-O from the Immcantation software suite^69, 70^. First, the FilterSeq.py function was used to remove reads with an average Q-score of less than 25. Then, MaskPrimers.py was used to identify the 14 base pair-long UMI appended during RT as well as the constant region isotype. The PairSeq.py function was used to retain only read pairs that passed both the FilterSeq.py and MaskPrimers.py processing steps. Consensus sequences for each UMI were then assembled using the BuildConsensus.py function, and the AssemblePairs.py function was used to assemble read mates into a single BCR, using sequential-guided assembly with an IMGT reference. The resulting sequences were then analyzed with IgBlast^72^ using reference sequences provided by IMGT. For comparison to single-cell BCR data, we used only unique, functional sequences supported by greater than 5 reads.

### Processing of cell hashing data

Cell hashing data was aligned to HTO barcodes using CITE-seq-Count v1.4.2. To establish thresholds for positivity for each HTO barcode, we first performed centered log-ratio normalization of the HTO matrix and then performed k-medoids clustering with k=5 (one for each HTO). This produced consistently five clusters, each dominated by one of the 5 barcodes. For each cluster, we first identified the HTO barcode that was dominant in that cluster. We then considered the threshold to be the lowest value for that HTO barcode among the cells classified in that cluster. To account for the scenario in which this value was substantially lower than the rest of the values in the cluster, we used Grubbs’ test to determine whether this threshold was statistically an outlier relative to the rest of the cluster. If the lower bound was determined to be an outlier at p=0.05, it was removed from the cluster, and the next lowest value was used as the new threshold. This procedure was iteratively applied until the lowest value in the cluster was no longer considered an outlier at p=0.05. Cells were then determined to be “positive” or “negative” for each HTO barcode based on these thresholds. Cells that were positive for multiple HTOs or were negative for all HTOs were excluded from downstream analysis. To account for differences in sequencing depth between samples, these steps were performed separately for each Seq-Well array that was processed.

## Data Availability

Raw data from this study has been deposited on GEO, accession number GSE232873.

## Supporting information

Supplemental Figures and Protocol

## Acknowledgements

We thank N. Kamelamela, C. Hallee, S. Levine, and the MIT BioMicro Center for assistance with library preparation and sequencing, as well as the KI Flow Cytometry Core and the New Iberia Research Center (NIRC).

## Author Contributions

Conceptualization: DMM, JCL, LC, IK

Methodology: DMM, YZ, JK, MM, SS, JL, NS, SUP

Investigation: DMM, YZ, JK, MM, SS, JL, NS, SUP

Visualization: DMM

Funding acquisition: JCL, LC, IK

Project administration: JCL, IK, LC

Supervision: JCL, IK, LC

Writing – original draft: DMM, JCL

Writing – review & editing: DMM, JCL, LM

