## Supplemental Figures and Protocol for "Full-length single-cell BCR sequencing paired with RNA sequencing reveals convergent responses to vaccination"

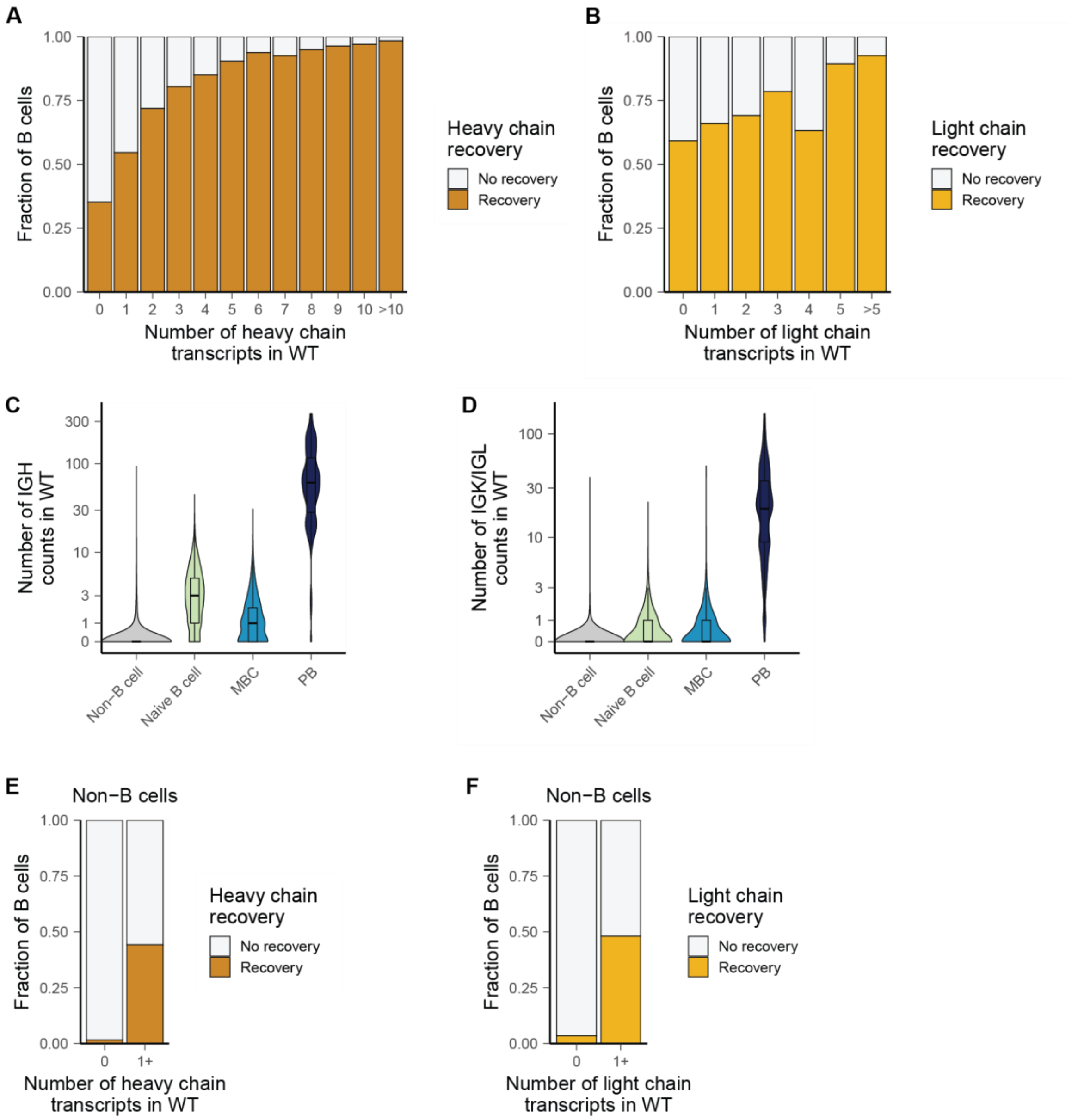


**Supplemental Figure 1**. A) Relationship between recovery of at least one functional BCR heavy chain from B cells sequence and the number of heavy chain transcripts detected in WT libraries. B) Relationship between the number of light chain transcripts detected in WT libraries of B cells and the recovery of at least one functional light chain sequence. C) Number of IGH (heavy chain) transcripts detected in single cells of each phenotype. D) Number of IGK/IGL (light chain) transcripts detected in single cells of each phenotype. E) Stacked bar plot of the recovery of heavy chain sequences from non-B cells and the number of heavy chain transcripts detected in WT libraries. F) Stacked bar plot of the recovery of light chain sequences from non-B cells and the number of light chain transcripts detected in WT libraries.

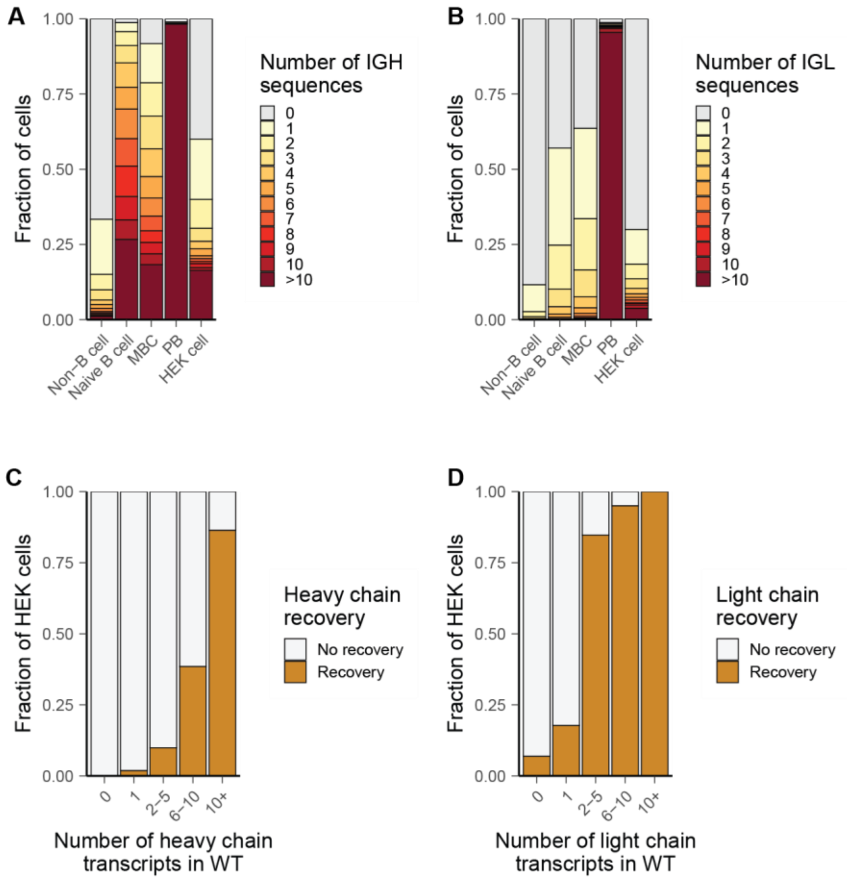


**Supplemental Figure 2**. A) Number of heavy chain molecules recovered in whole transcriptome libraries from each phenotype in HEK cell experiment. B) Number of light chain molecules recovered in whole transcriptome libraries from each phenotype in HEK cell experiment. C) Relationship between heavy chain recovery and number of heavy chain transcripts in WT libraries among HEK cells. D) Relationship between light chain recovery and number of light chain transcripts in WT libraries among HEK cells.

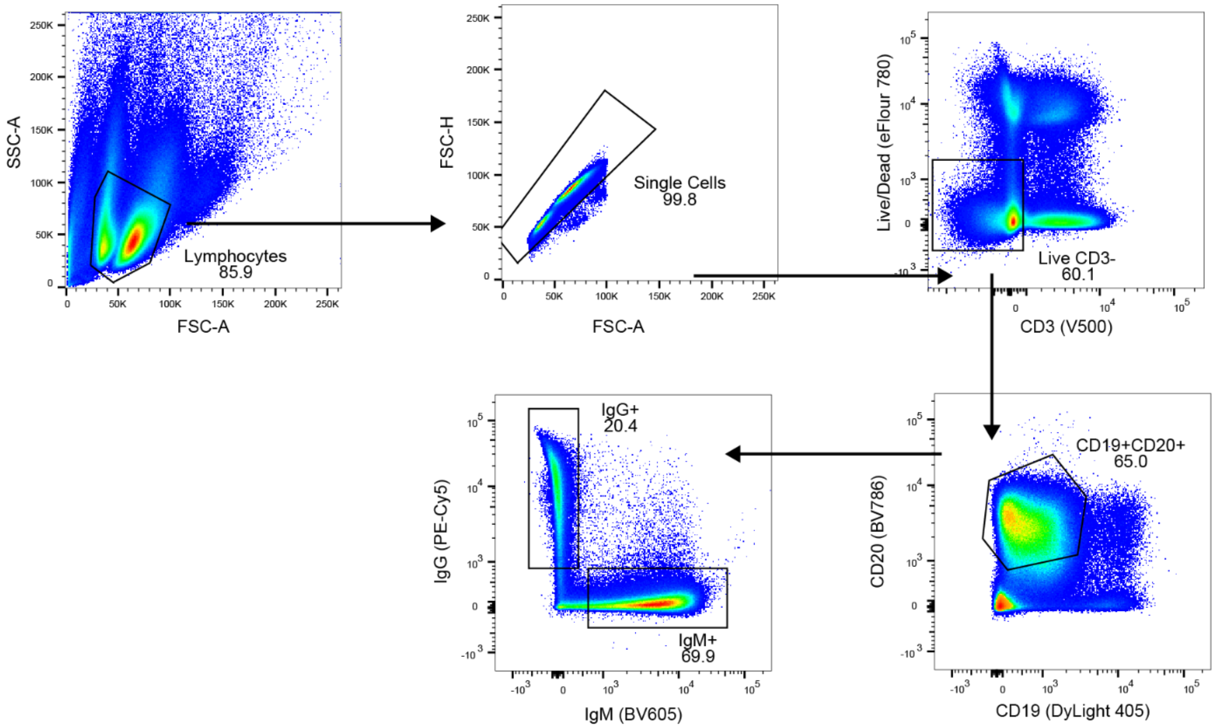


**Supplemental Figure 3.** Flow cytometry gating strategy. B cells were gated as follows: lymphocytes -> single cells -> live CD3- -> CD19+CD20+, and then as either IgG+ or IgM+.

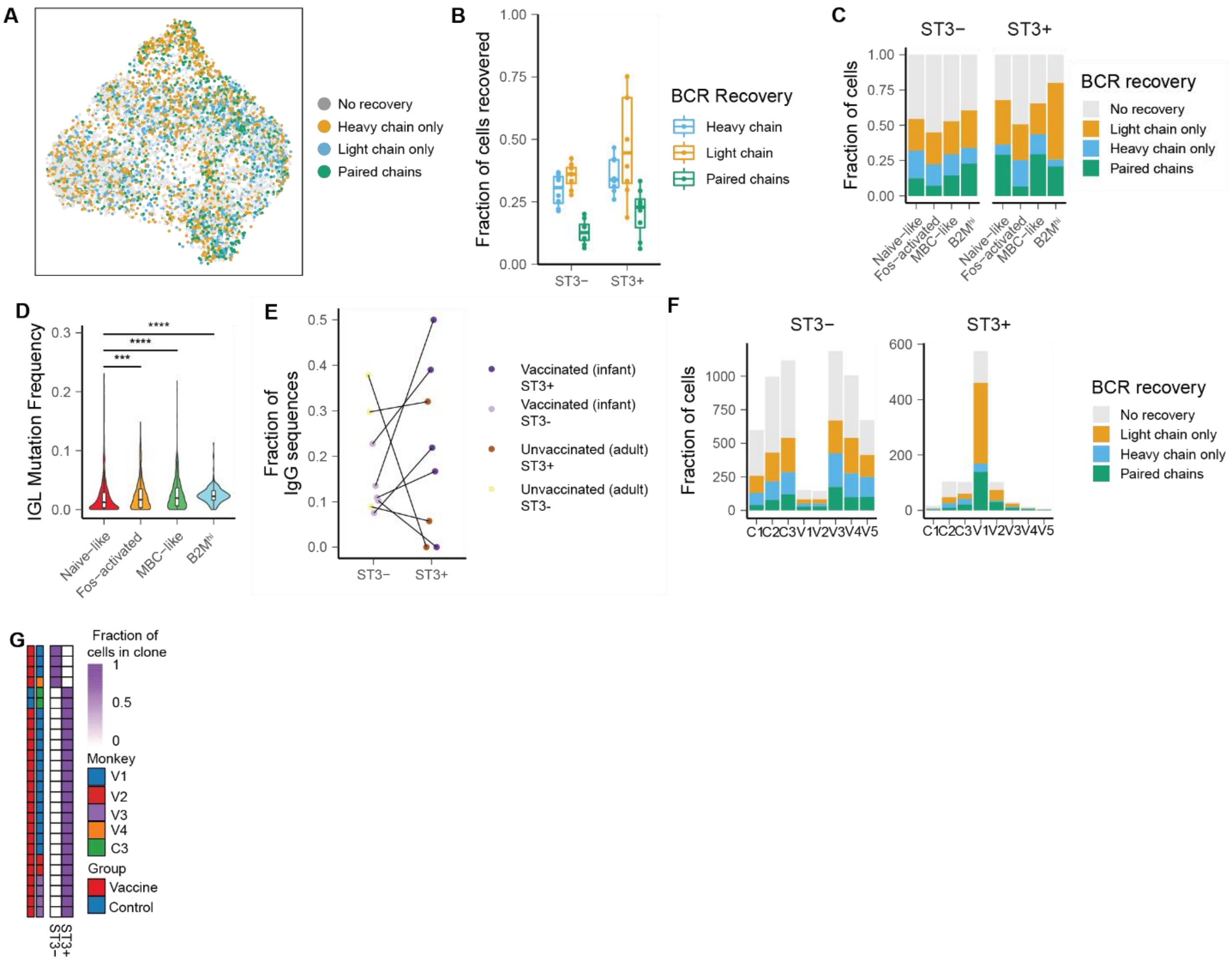


**Supplemental Figure 4.** A, B, C) Recovery of BCR sequences from single cells. D)Frequency of somatic mutation in light chain sequences recovered from each phenotype. P-values are calculated with a two-sided Wilcoxon rank-sum test and are adjusted with Bonferroni correction. E). Fraction of IgG sequences among ST3+ and ST3- cells from each monkey. F) Number of cells with BCR recovery from each monkey and sort fraction. G) Fraction of cells from each expanded BCR clonotype recovered from ST3- and ST3+ cells from each sort fraction.

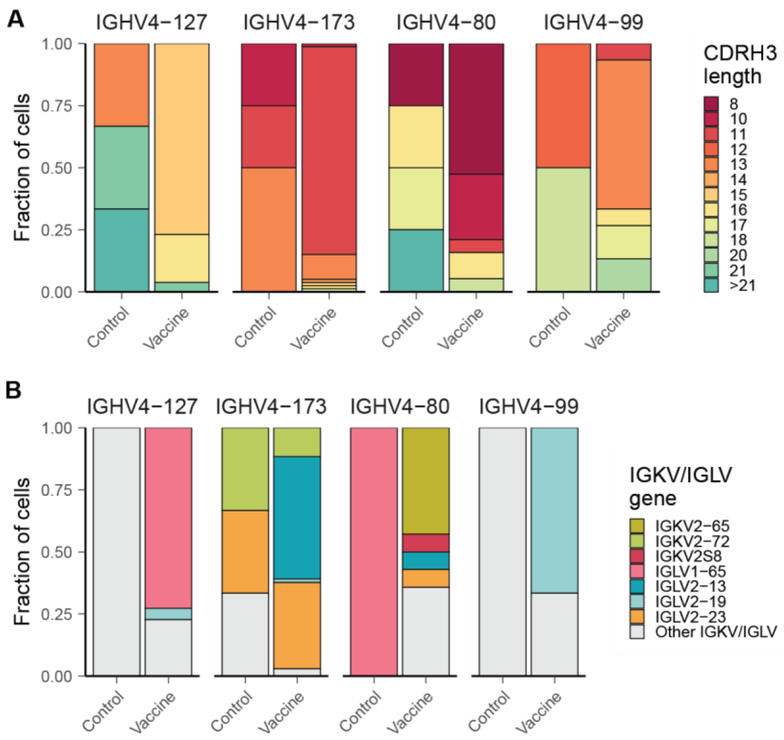


**Supplemental Figure 5**. A) CDRH3 lengths of ST3-reactive BCR using IGHV genes statistically enriched among ST3-reactive cells in vaccinated monkeys, separated by vaccinated (infant) and unvaccinated (adult) monkeys. B) Light chain pairings of ST3-reactive BCR using genes statistically enriched among ST3-reactive cells in vaccinated monkeys, separated by vaccinated (infant) and unvaccinated (adult) monkeys.

Supplemental Table 3: Expanded ST3-reactive clonotypes recovered from unvaccinated adult monkeys.

| **Monkey** | **Phenotype** | **IGHV** | **CDRH3** | **Isotype** | **IG(K/L)V** | **Mutation Frequency** |
|---|---|---|---|---|---|---|
| C3 | MBC-like | IGHV3S42*01 | YYCADGRLAL | IgM | NA | 0.129 |
| C3 | MBC-like | IGHV3S42*01 | YYCADGRLAL | IgM | IGLV1S4*01 | 0.129 |
| C3 | MBC-like | IGHV3-28*02 | CARDPGGYYCSW | IgD | NA | 0.041 |
| C3 | MBC-like | IGHV3-28*02 | CARDPGGYYCSW | IgD | NA | 0.041 |

**Supplemental Protocol**

This protocol assumes that whole transcriptome products generated by Seq-Well or Dropseq are being used.

**Materials**:

Biological Materials

- 3'-barcoded whole transcriptome amplified products generated by the SeqWell or DropSeq platform.

Reagents

- 96 rxn xGen® Lockdown® Reagents (IDT, cat. no. 1072281)
- Pull-down-probes (IDT, see **Supplementary Table** for oligo sequences)
- Human Cot-1 DNA (Invitrogen, cat. no. 15279011)
- Dynabeads™ M-270 Streptavidin (Invitrogen, cat. no. 65306)
- KAPA HiFi HotStart ReadyMix (Roche, cat. no. KK2602)
- DNAse, RNAse free UltraPure Distilled Water (Invitrogen, cat. no. 10977-015)
- Sera-mag SpeedBeads (Fisher, cat. no. 09-981-123)
- PEGN8000 (Amresco, cat. no. 0159)
- 0.5 M EDTA, pH 8.0 (Amresco, cat. no. E177)
- 1.0 M TrisNHCL, pH 8.0 (Amresco, cat. no. E199)
- Tween 20 (Amresco, cat. no. 0777)
- 5 M NaCl (VWR, cat. no. 97062-858)
- 200-proof pure Ethanol (Koptec, cat. no. V1001)

Definition of terms

- WTA: Whole transcriptome amplification product. Full-length transcriptome products that are single-cell barcoded on the 3' side. In this protocol, WTA libraries were constructed via Seq-Well, which has an identical structure to DropSeq.
- Pull-down probes: Oligonucleotide probes with biotinylated 5' end. This protocol uses probes targeting the constant region of the BCRs. Used in conjunction with the IDT xGen® Lockdown® Reagents kit to target BCR whole transcripts. See Supplemental Table 4 for exact sequences.
- UPS primers: Universal priming site primers. UPS and UPS2 primers are used to uniformly amplify all products and construct Illumina-compatible libraries. See Supplemental Table 4 for exact sequences.

**Procedures**

I. Preparation
   1. Thaw xGen® Lockdown® Reagents and make proper dilutions. Wash buffer 1 can be brought to 37˚C before use to make sure all salts are dissolved.
   2. Pull-down probes should be diluted ahead of time. We recommend aliquoting the probes into a multi-channel compatible format such as in strip tubes with caps. (See Supplemental Table 4 for exact oligo sequences)
   3. IGH probes should be combined and diluted to 1.5 uM each.
   4. IGK and IGL probes should be combined and diluted to 1.5 uM each.

II. BCR enrichment: hybridization to pull-down probes
   1. Make the following master mix. Make two separate mixes for each sample/reaction (one for IGH and the other for IGK+IGL).

| | |
|---|---|
| UPS primer (50 uM) | 0.8 uL |
| Human Cot-1 DNA (Invitrogen, cat. no. 15279011) | 0.5 uL |
| xGen 2x Hybridization Buffer (IDT, cat. no. 1072281) | 8.5 uL |
| xGen Buffer Enhancer (IDT, cat. no. 1072281) | 2.7 uL |
| Total | 12.5 uL |

2. Mix 3.5 uL of WTA product with the master mix to make up a reaction volume of 17 uL.
3. Incubate mixture at room temperature for 5 min.
4. Incubate the mixture at 95 ˚C for 10 min (on thermocycler).
5. Quickly add 1 uL of IGH or IGK+IGL Pull-down probe cocktail
6. Incubate the mixture at 65 ˚C for 60 min to allow for hybridization of pull-down probes to the WTA libraries.

III. BCR enrichment: prepare the pull-down Streptavidin beads (Invitrogen, cat. no. 65306)

1. Aliquot 50uL of beads per sample into a 1.5 mL microcentrifuge tube.
2. Place the tube on a magnetic stand and allow for the beads to pellet.
3. Remove the supernatant and add an equal volume of bead wash buffer into the tube.
4. Vortex to mix, and place back on the magnetic stand
5. Repeat 3.3 – 3.4
6. Take the tube off of the magnet, and add an equal volume of bead wash buffer
7. Aliquot 50uL (per sample) of the mixture into PCR strip tubes.
8. Place tubes back on magnetic stands.
9. Remove the supernatant. The beads are now ready for hybridized mixes from II.

IV. BCR enrichment: prepare wash buffers (during III)

1. Prepare 300 uL/sample Wash Buffer 1 (WB1) by 1:10 dilution (IDT, cat. no. 1072281). Split 300 uL/sample WB1 into 100 uL/sample and 200 uL/sample.
2. Prepare 400 uL/sample Stringent Wash Buffer (SWB) by 1:10 dilution (IDT, cat. no. 1072281). Aliquot 400 uL/sample SWB into 200 uL + 200 uL/sample.
3. Prepare 200 uL/sample of each Wash Buffer 2 (WB2) and Wash Buffer 3 (WB3) by 1:10 dilution (IDT, cat. no. 1072281).

V. BCR enrichment: Pull-down BCR transcripts from WTA libraries (after pull-down probes have been hybridized to WTA libraries).

1. Add hybridized mixture from II into prepared beads from III.
2. Gently vortex to mix
3. Incubate mixture at 65˚C for 45 min
    - 3.1. Preheat 100 uL/sample of WB1 and 400uL/sample of SWB (2x200uL) during incubation of 5.3. We recommend aliquot the buffers into PCR strip tubes and preheat on the thermocycler or heated mixer.
    - 3.2. Intermittently vortex the mixture every 10 minutes to keep the streptavidin beads suspended in the mixture
    - 3.3. If possible, we recommend using a thermal shaker to automate intermittent vortexing (e.g. Eppendorf ThermoMixer C)
4. After incubation, add 100uL of heated WB1 into each hybridized mixture and place onto a magnetic stand
5. Aspirate supernatant, and take mixtures off of the magnetic stand.
6. Add 200uL of heated SWB into each reaction, and resuspend the bead pellets
7. Incubate mixture at 65 degrees for 5 minutes
8. Place mixture back on the magnetic stand, and repeat 5.5 – 5.7
9. Aspirate the final wash of SWB
10. Add 200uL of room temperature WB1 into each reaction
11. Vortex for 2 minutes to mix, spin to collect liquid, and place on the magnetic stand.

12. Aspirate, and add 200uL of room temperature WB2.
13. Vortex for 1 minute to mix, spin to collect liquid, and place on the magnetic stand.
14. Aspirate, and add 200uL of room temperature WB3.
15. Vortex for 30 seconds to mix, spin to collect liquid, and place on the magnetic stand.
16. Aspirate, and add 20uL of water.

VI. BCR enrichment: PCR amplify pull-down BCR products

1. For each sample/reaction, perform 5 PCR reactions using 2 uL of the mixture from V (using a total of 10 uL out of 20 uL). Make the following master mix per sample/reaction:

| | |
|---|---|
| UPS primer (10 uM) | 2.0 uL |
| 2x Kapa Hifi Hotstart Readymix (Roche, cat. no. KK2602) | 12.5 uL |
| DNAse-free water (Invitrogen, cat. no. 10977-015) | 8.5 uL |
| Total | 23 uL |

2. Aliquot 5 x 23 uL of master mix for each sample/reaction
3. Add 2 uL of the mixture from V into each of the aliquots
4. PCR amplification using the following condition

1 cycle of
[95˚C for 3 minutes]
25 cycles of
[98˚C for 40 seconds
67˚C for 20 seconds
72˚C for 1 minute]
1 cycle of
[72˚C for 5 minutes]
Hold at 4˚C

5. After amplification, pool all 5 PCR reactions (20uL for each sample) into a single tube (for 100uL total)
6. Perform 0.65x SPRI for IGH or 0.8x purification for IGL/IGK (see appendix for detailed SPRI protocol)
7. Elute the samples into 13-15 uL of DNAse-free water
    - We recommend assessing the quality of the enrichment via a fragment analyzer
    - If amplification was unsuccessful (either due to low concentration of the product, or technical error), the rest of the beads from V can be used to repeat VI

VII. BCR V-region capture and extension:

1. Prepare BCRV-UPS primer cocktails for heavy chain and light chain separately. Equimolar pool each of the primers. Dilute as necessary to 10uM (total concentration). (See Supplemental Table 4 for exact oligo sequences)
2. Make the following master mix per sample/reaction. Make two separate mixes (one for IGH and the other for IGK+IGL)

| | |
|---|---|
| BCRV-UPS2 primer cocktail (10 uM) | 2.5 uL |
| 2x Kapa Hifi Hotstart Readymix (Roche, cat. no. KK2602) | 12.5 uL |
| DNAse free water (Invitrogen, cat. no. 10977-015) | 6.0 uL |
| Total | 21 uL |

3. Aliquot 21 uL per PCR reaction of master mix into PCR tubes or plates
4. Add 4 uL of BCR-enriched product for a total volume of 25 uL per PCR reaction

5. Perform V-region capture and extension using the following condition:

1 cycle of
[95˚C for 5 minutes]
1 cycle of
[55˚C for 30 seconds]
1 cycle of
[72˚C for 2 minutes]
Hold at 4˚C

6. After extension, add 25 uL of DNAse-free water and bring each reaction to a total volume of 50 uL
7. Perform 0.65x SPRI for IGH or 0.8x purification for IGL/IGK (see appendix for detailed SPRI protocol)
8. Elute the samples into 11 uL of DNAse-free water

VIII. BCR UPS2 amplification

1. Prepare UPS2-N7xx and UPS-mod-N5xx primers to amplify the resulting BCR products and add Illumina flanking sequences for sequencing. Thaw and dilute the primers into 10 uM aliquots. (See Supplemental Table 4 for exact oligo sequences)
2. For each sample/reaction, perform 4 PCR reactions. Make the following master mix for each sample/reaction:

| | |
|---|---|
| 2x Kapa Hifi Hotstart Readymix (Roche, cat. no. KK2602) | 12.5 uL |
| UPS-mod-N5xx (10 uM) | 0.5 uL |
| UPS2-N7xx (10 uM) | 0.5 uL |
| DNAse free water (Invitrogen, cat. no. 10977-015) | 9.0 uL |
| Total | 22.5 uL |

3. Aliquot 4 x 22.5 uL of master mix for each sample/reaction
4. Add 2.5 uL of products from VII for a total volume of 25 uL per PCR reaction
5. PCR amplification using the following conditions:

1 cycle of
[95˚C for 2 minutes]
10-18 cycles of
[95˚C for 30 seconds
60˚C for 30 seconds
72˚C for 1.5 minutes]
1 cycle of
[72˚C for 5 minutes]
Hold at 4˚C

6. Take 12.5 uL of each reaction and pool a total volume of 50 uL per sample/reaction
7. Perform 0.65x SPRI for IGH or 0.8x purification for IGL/IGK (see appendix for detailed SPRI protocol)
8. Elute the samples into 13 uL of DNAse-free water

IX. Sequencing of BCR libraries

- We recommend sequencing on the Illumina MiSeq using the 600-cycle kit or NovaSeq S1 using the 500-cycle kit
- The sequencing specification is shown below. See Supplemental Table 4 for detailed sequence primer sequences.

| | Sequencing Primer | MiSeq | NovaSeq |
|---|---|---|---|
| Read 1 | SeqWell/DropSeq Read 1 Seq Primer | 20 bp | 20 bp |
| Index Read 1 | IGH, IGK, IGL Seq Primers | 300 bp | 252 bp |
| Index Read 2 | Nextera Index Read 2 Seq Primers | ___ | 8 bp |
| Read 2 | Nextera Read 2 Seq Primers | 300 bp | 252 bp |

Appendix. SPRI protocol

I. Making SPRI reagent
   1. In a 50 mL conical using sterile stock solutions, prepare 1X TE by 1:10 dilution from 10X TE buffer solution (Teknova, cat. no. T3457).
   2. Mix Sera-mag SpeedBeads well and transfer 1 mL to a 1.5 mL microtube.
   3. Place SpeedBeads on a magnet stand until beads are drawn to the magnet.
   4. Remove supernatant with P200 or P1000 pipetter.
   5. Add 1 mL TE to beads, remove from the magnet, mix, and return to the magnet.
   6. Remove supernatant with P200 or P1000 pipetter.
   7. Repeat 5 – 6
   8. Add 1 mL TE to beads and remove from the magnet. Fully resuspend.
   9. Add 9 g PEGN8000 to a new 50 mL sterile conical tube.
   10. Add 10 mL 5 M NaCL (or 2.92 g) to the conical.
   11. Add 500 µL 1 M TrisNHCL to the conical tube.
   12. Add 100 µL 0.5 M EDTA to the conical.
   13. Fill conical to ~ 48 mL using sterile dH20. You can do this by eye, just go slowly.
   14. Mix conical for about 15 minutes until PEG goes into solution (solution, upon sitting, should be clear).
   15. Add 275 uL 10% Tween 20 to the conical and mix gently.
   16. Mix 1 mL SpeedBead in TE solution and transfer to 50 mL conical.
   17. Fill conical to 50 mL mark with dH20 (if not already there) and gently mix 50 mL conical until brown.

II. SPRI clean up
   1. Prepare fresh aliquots of 80% EtOH
   2. Mix the appropriate volume of SPRI reagent with the sample DNA library
   3. Incubate mixture for 10 min at room temperature
   4. Place on a magnet stand and wait for 2 min
   5. Remove supernatant
   6. Add 180 uL 80% EtOH
   7. Mix by moving the entire tube or plate across the magnet stand so that the beads migrate through the EtOH solution approximately 6 times
   8. Remove and discard EtOH without touching the beads
   9. Repeat steps 6 – 8
   10. Allow the bead pellet to dry. Roughly 5 – 10 min
   11. Elute in an appropriate volume of water
   12. Incubate at room temperature for 10 min
   13. Place back on the magnetic stand for beads to pellet
   14. Collect the supernatant and discard the magnetic beads.
